# BLASE: Bulk Linkage Analysis for Single Cell Experiments - Teasing Out the Secrets of Bulk Transcriptomics with Trajectory Analysis

**DOI:** 10.1101/2025.09.03.673925

**Authors:** Andrew McCluskey, Toby Kettlewell, Adrian M. Smith, Rhiannon Kundu, David A. Gunn, Thomas D. Otto

**Affiliations:** School of Infection and Immunity, University of Glasgow, Glasgow, UK; School of Mathematics and Statistics, University of Glasgow, Glasgow, UK; Unilever R&D, Colworth Science Park, Bedford, UK; Unilever R&D, King’s College London, London, UK; LPHI, CNRS, INSERM, Université de Montpellier, Montpellier, France; Bernhard Nocht Institute for Tropical Medicine, Hamburg, Germany; University of Hamburg, Hamburg, Germany

## Abstract

1

**Motivation:** scRNA-seq experiments can capture cell process trajectories. Bulk RNA-seq is more practical, however does not have the granularity to elucidate cell-type specific trajectories. Deconvolution methods can estimate cell-types in RNA-seq data, but there is a need for methods characterising their pseudotime.

**Results:** We show that our method, BLASE, can identify the progress of an RNA-seq sample through a trajectory in a scRNA-seq reference.

**Conclusion:** BLASE can be used to a) annotate scRNA-seq data from existing RNA-seq, b) identify progress of RNA-seq data through a process based on scRNA-seq data, and c) be used to correct developmental differences in RNA-seq differential expression analysis.

## 2 Background

Transcriptomics revolutionised our understanding of cells in disease and health [1, 2, 3, 4]. In single-cell transcriptomic sequencing (scRNA-seq), the analysed dataset may contain information about cell types (or subtypes), and a biological process (calculated by “trajectory inference” methods, represented as “pseudotime,” a continuous measure of the progress of a cell through the process). Several tools (such as CIBERSORTx [5]) have successfully shown that scRNA-seq can be used to deconvolute cell types within a bulk RNA-seq sample, providing insights at a greater granularity than analysis that looks at bulk data in isolation. These methods typically make use of a scRNA-seq reference to decompose the cell types that constitute a bulk RNA-seq sample into proportions. These tools allow the user to reuse existing datasets, which may be hard to reproduce with scRNA-seq if originally generated from rare or valuable samples. Researchers may also wish to use deconvolution to reduce cost, as in some cases, bulk RNA-seq with deconvolution can provide sufficient insight into the biology, without needing to use costly and challenging scRNA-seq methods. In recent reviews [6, 7], DWLS and MuSiC [8, 9] were both assessed as being highly accurate. These methods are used with already annotated scRNA-seq reference datasets. CIBERSORTx generates “signature matrices” based on the differential expression of genes in each annotated cell type in the reference. Using these matrices, CIBERSORTx then uses Support Vector Regression (SVR) to estimate proportions of the cell types present in a bulk sample. DWLS (Dampened Weighted Least Squares) also generates a signature matrix, based on marker genes from differential expression analysis. DWLS uses a WLS method on each bulk sample, and then uses a dampening constant to prevent overweighting of rare or low expression cell types. MuSiC (Multi-Subject Single Cell deconvolution) also uses a reference scRNA-seq dataset to calculate signature matrices for each annotated cell type. It then uses a Weighted Non-Negative Least Squares (W-NNLS), where each gene is weighted (to give greater weight to informative genes), to estimate cell type proportions in a bulk sample.

These current deconvolution methods focus primarily on identifying the proportions of cell types contained within a bulk RNA-seq sample. However, it can be desirable to identify how far a bulk sample has progressed through a biological process. Because of the nature of these processes, the differences are typically gradual changes over pseudotime, as opposed to the discrete differences between mature cell types. Thus, for two points close in pseudotime, differences can be limited and different considerations are required to separate those points in terms of gene expression. This leads to cell-type deconvolution techniques giving unrealistic results (such as estimating that pseudobulks from multiple windows over pseudotime are mostly comprised of cells from the first pseudotime bin) when used to identify the position of a bulk sample in pseudotime.

We postulated that it should be possible to decompose bulk RNA-seq of a process (e.g. disease changes, cell development in the bone marrow, or pathogen lifecycle) using existing scRNA-seq data. Trajectory inference is a popular method for exploring biological processes in scRNA-seq data, with a variety of methods proposed (Slingshot, Destiny, Palantir, Monocle [10, 11, 12, 13]). These methods assign a numerical value to each cell in a dataset, indicating their progress through a biological process of interest. Importantly, pseudotime is a continuous variable, whereas cell type is typically considered in discrete terms. Methods have been developed to complement trajectory inference by calculating which genes are significantly associated with pseudotime (for example TradeSeq’s `associationTest` [14]), and whether genes are differentially expressed between two conditions in the same process (for example, TradeSeq’s `conditionTest`, PseudotimeDE [15], and TrAGEDy [16]). There is an additional difference in the matter of identifying the position of a bulk sample in pseudotime compared to cell-type deconvolution; namely, that only a single point in the process should be the expected result for a well synchronised sample (i.e. a sample where every cell is at a similar point in pseudotime), as opposed to the proportions given by cell-type deconvolution methods, which makes benchmarking challenging. Because of these, we reasoned that cell-type deconvolution methods are inappropriate for bulk pseudotime deconvolution.

In such a tool for identifying the progress of a bulk through a biological process, we would desire a clear mapping to a specific time-point, with a measure of confidence, and the flexibility to use this information to correct for differences in pseudotime between conditions. Interestingly, so far, only one option exists for the deconvolution of bulk RNA-seq over a trajectory: MeDuSA (mixed model-based deconvolution of cell-state abundances). MeDuSA is similar to other deconvolution tools in that it predicts proportions, but is adapted for use on trajectories. MeDuSA uses a mixture model to deconvolute the abundance of cells over a one dimensional trajectory, estimating the distribution of cells over pseudotime. However, in our testing, MeDuSA could not always resolve the correct distribution of simulated samples based on a scRNA-seq of *Caenorhabditis elegans* used in their paper (Figure A.1) or synchronised *Plasmodium falciparum* bulk RNA-seq samples well. Instead, it overestimated the period of pseudotime through which a bulk was distributed. As MeDuSA and other cell-type deconvolution tools do not accommodate all of these desirable features, we decided to develop BLASE.

Here, we present this novel tool in an open-source R package that provides a toolkit for bulk pseudotime identification. BLASE uses a correlation-based method to find the most similar interval of a discretised pseudotime in a scRNA-seq dataset to a bulk RNA-seq sample. We demonstrate that BLASE can accurately map pseudobulked simulated and genuine scRNA-seq data and outperform existing tools. Using published real data, we deconvoluted disease progress of psoriasis in spatial data, and find and correct a growth-rate difference in a parasite mutant.

## 3 Results

Here we present BLASE, which maps bulk RNA-seq data onto a scRNA-seq trajectory (Figure 1), which will be publicly available through Bioconductor as an R package. We show that current cell-type deconvolution methods do not perform well on trajectories to motivate our implementation. Finally, we share three use cases showing how BLASE can be used on real data.

**Figure 1.**
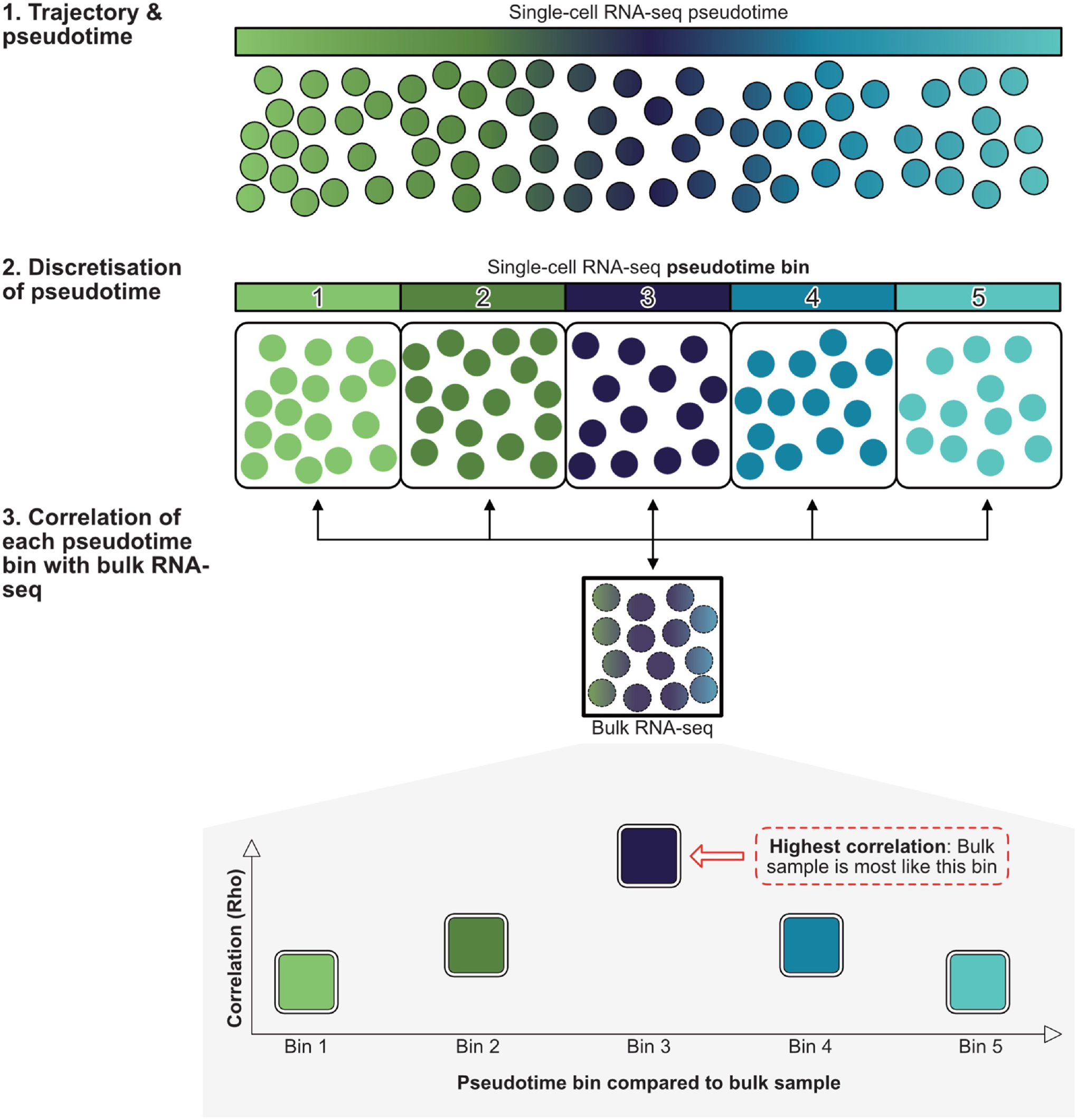
A schematic of the BLASE algorithm for identifying the progress of a bulk RNA-seq sample through a known scRNA-seq trajectory. Initially, a single-cell reference dataset with trajectory data is used to define discrete “pseudotime bins”. Next a bulk sample is compared against each bin, using the the Spearman’s Rho of their normalised counts. The pseudotime bin with the highest Rho is considered a match.

### 3.1 Validation 1: BLASE outperforms existing tools on simulated data

To validate BLASE on a dataset with a known ground truth, we generated a simulated dataset with Dyngen [17]. Pseudotime and a trajectory were calculated by Slingshot [10], and cells were grouped into 5 bins by pseudotime range. For each bin we generated two replicate pseudobulks and used BLASE and other deconvolution tools to “decompose” them against the pseudotime. BLASE generated the expected outcome (Figure A.2) of aligning each pseudobulk sample to its correct pseudotime bin. BLASE calculated a high correlation, showing that the rank-order correlation of genes is high, and indicates these are confident results, which means that bootstrapping has shown that the predicted bin is likely to have the highest true correlation (see Methods).

To compare our results with existing tools, we used CIBERSORTx, DWLS, MeDuSA, and MuSiC to decompose the same pseudobulks (Figure A.2). These four existing tools returned the expected proportions of cell categorisations, rather than a single best mapping score. In order to compare them with BLASE, we considered the cell-type predicted to have the highest proportion to be the call made by a given tool. CIBERSORTx was unable to perform as expected on this dataset despite various troubleshooting and was excluded from the analysis. DWLS and MuSiC correctly assigned the bins. DWLS mapped a mean of over 91% of cells to the correct bins, whereas MuSiC had a mean correct mapping of 84.2%. MeDuSA generally gave the highest proportion to the correct bin, but only assigned a mean of 42.3% of cells to the correct bin, showing a much wider distribution than was expected (Table A.1).

Overall, this shows that BLASE is not only accurate, but also that it can match the performance of leading cell type decomposition methods for this simulated data.

### 3.2 Validation 2: Decomposition of myeloid cells in haematopoietic lineages

The haematopoietic niche is located in bone marrow, where new immune cells differentiate from pluripotent stem cells into highly specialised cells which are released into the blood. This transformation is a continuous process, and thus this is a perfect example to test and compare BLASE on with real data.

First, we took the myeloid cell lineage from Mende et al. 2020 [18] (Figure 2 a, black box). Next, we projected those cells into a PHATE embedding, calculated pseudotime with Slingshot and created 5 bins with equal pseudotime lengths (see subsection 6.4, similar to the classes of cell selected in a.

**Figure 2.**
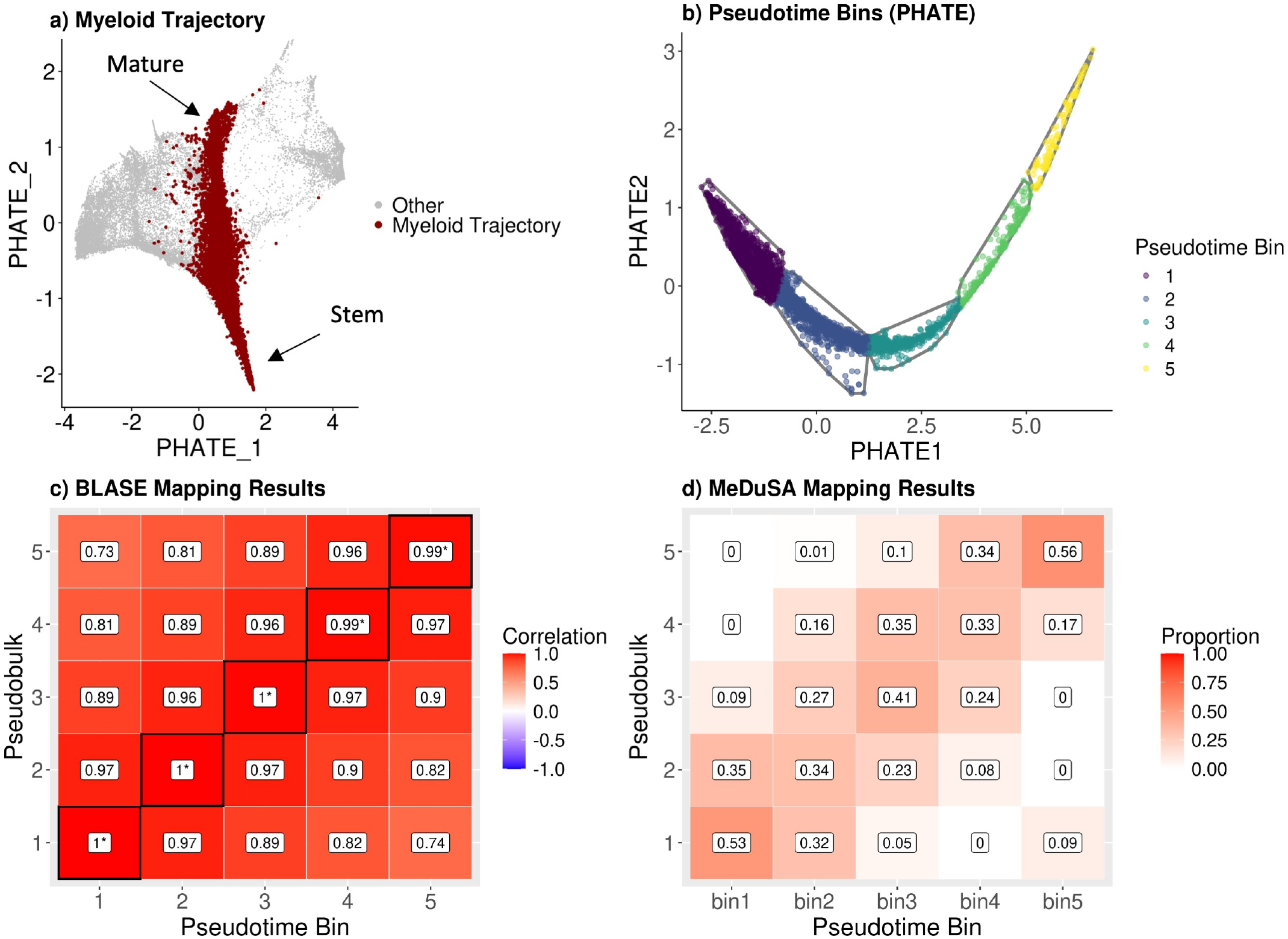
Validating BLASE with a test/train pseudobulked dataset of haematopoiesis. a, The myeloid cell differentiation lineage selected for testing BLASE results, highlighted on all other cells in the PHATE projection. b, A PHATE embedded plot of the single-cell training dataset (including only the myeloid differentiation lineage) coloured by assigned pseudotime bins. c, A heatmap showing Spearman’s Rho (of the gene list calculated by TradeSeq) between pseudotime bins and bulk RNA-seq samples. Asterisks indicate confident results, and black outlines indicate the best match. Pseudotime bins are shown on the x-axis, tested bulk samples are on the y-axis. d, The mapping proportions from MeDuSA deconvoluting the same data as c.

with BLASE (Figure 2 b). To create a bulk query dataset, we split the dataset in half at random, and then pseudobulked one half by the pseudotime bin (Figure A.3). The non-pseudobulked half was used a reference scRNA-seq dataset.

We then applied BLASE and other cell type deconvolution methods (see Figure A.2 for a comparison) to predict which pseudotime bin had been the origin of each pseudobulk in the myeloid lineage. BLASE correctly and confidently maps each pseudobulked bin to the correct reference bin, showing that BLASE is capable of overcoming the intricacies of scRNA-seq data, as shown by the heatmap in Figure 2 c. This heatmap shows the Spearman correlation as calculated by BLASE for each pseudobulk to each reference pseudotime bin. The highest correlation as predicted by BLASE is shown by an outlined tile, and asterisks indicate “confident” results, where BLASE has calculated that it is confident that a bin other than the best mapped one is not the best match. The heatmap shows a high correlation for other bins in addition to the correct one. This is expected, as over a continuous process, we expect differences in expression to be smooth over pseudotime. We tested several gene lists with BLASE, and found that using TradeSeq, gene peakedness, and gene peakedness spread all gave completely correct and confident results, but doing so with every available gene did not (see subsubsection 3.4.2). We believe this is because only some genes are relevant to this process, and using all genes introduces noise that obscures the true transcriptomic signature of myeloid differentiation.

It is noteworthy that the other tested cell-type deconvolution tools performed less well at finding the true paired bin than BLASE when provided the same training/test data. DWLS predicted that every pseudobulked bin should map to the first bin of the reference (Figure A.2, b). This resulted in DWLS mapping a mean of only 20.9% of cells correctly. CIBERSORTx called the start and end bulk samples well, however it performed poorly on the pseudobulk samples which should have mapped to bins 2 and 3, resulting in a mean correct mapping of only 46.3%. MuSiC (mean correct mapping of 75.0%) maps the bulks well, without the smoothing of the pseudotime as observed with BLASE. However, the proportion of mapped cells in bin 4 drops from the expected 100% to just 56%. It is important to remember that these other tools were designed for use with discrete cell types and not the continuum of pseudotime. Medusa, on the other hand, accounts for this continuous process. It captures the smooth distribution, as an “association” of the bins with the bulk data, however it only correctly maps a mean of 43.3% of cells (Figure 2, d). BLASE however, finds an optimum and uses bootstrapping to identify high confidence results. Overall, three tools can accurately map the pseudobulks onto the developmental process of this myeloid cell data, however only BLASE can map the bulk sample with statistical confidence to one time point.

### 3.3 Use Case 1: BLASE resolves keratinocyte differentiation in the epidermis in spatial data

Keratinocytes are a key cell type in the skin and constitute a major part of the environmental barrier that the tissue provides. These cells undergo a range of transcriptional changes in the epidermis as they develop. Basal keratinocytes are stem cell like, and proliferative. Some will commit to differentiation, detaching from the base of the epidermis, and building the proteins required for skin function, taking on a spiny appearance, thus “spinous” keratinocytes. These then develop into granular keratinocytes which continue producing substances that are necessary for epidermal function, including keratohyalin granules, which are required for the formation of the cornified envelope. Following this, the keratinocytes develop into enucleated corneocytes, which are effectively dead, but form the final barrier of skin [19]. This process can be captured with Spatial Transcriptomics (ST). In the dataset generated by Ma et al. in 2023 [20], the authors use 10x Visium to investigate psoriasis at a spatial resolution. The 10x Visium platform works on a matrix of “spots”, over which tissue is arranged. The cells in each spot are then barcoded and sequenced together. This produces count data for each spot, as the cells in a given spot are sequenced, and their spatial position is known based on the id of the spot. This results in a dataset which is formed of over a thousand spots, each effectively a very small bulk RNA-seq sample of the few cells contained in the spot. Using BLASE, we can then attempt to decompose where the cells in each spot belong on the axis of pseudotime, revealing more about the spatial characteristics of cellular processes, in this case, keratinocyte differentiation.

First we built the pseudotime from a dataset of neonatal epidermal cells, which we subset to include only keratinocytes [21] (Figure 3, a). This trajectory covered the transition from basal to spinous and, finally, granular keratinocytes (corneocytes no longer produce RNA, and so are not detected by transcriptomics). These cell types were identified using markers defined in the original paper (Basal: KRT14, KRT5, Spinous: KRT1, KRT10, Granular: DSC1, KRT2) see Figure 3, b-d. These are the key cell states as the keratinocytes differentiate and travel upward to the outer layers of the epidermis. Next we mapped the different spots of the 10x Visium to the pseudotime, predicted the best matching bin and projected it back onto the spatial map (Figure 3). One of the symptoms of psoriasis is an over-abundance of keratinocytes, forming psoriatic lesions. By applying BLASE to these samples, we can see a strong prediction of developed granular keratinocytes in a thicker epidermal layer (see the histology image in Figure 3, e), compared with the dermal layer, where few keratinocytes would be expected. By localising these known cell processes in ST data, new understanding of spatio-temporal relationships to transcriptomic processes may be possible.

**Figure 3.**
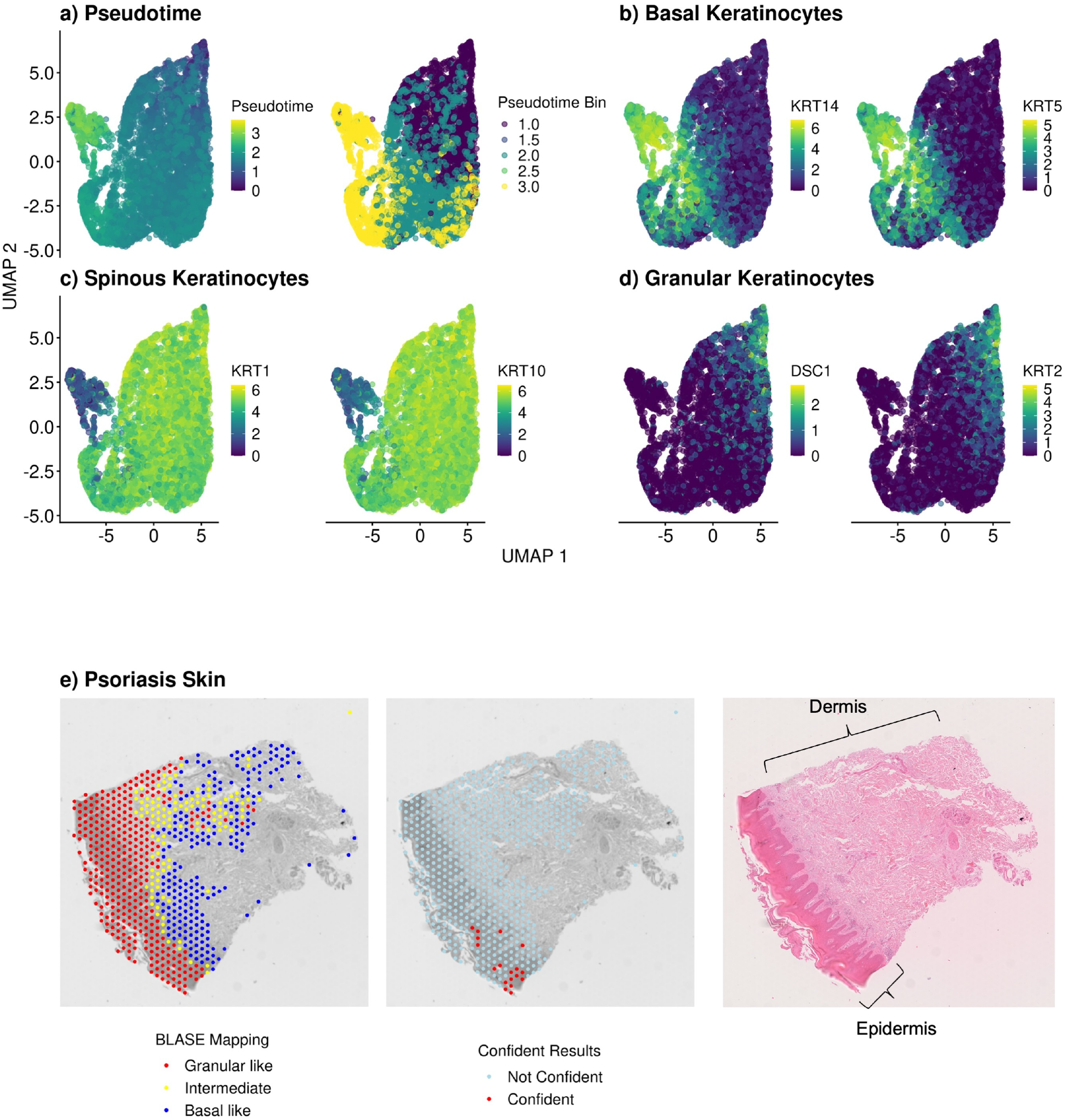
Temporal deconvolution of Visium spots. a, Pseudotime calculated by Destiny and the five pseudotime bins calculated by BLASE in the reference. b-d, Expression of two marker genes of Basal (b), Spinous (c), and Granular (d) Keratinocytes in the reference. e, Estimated progression mappings and confidence from BLASE, and a histology image (haematoxylin and eosin (H&E) staining) for a sample of Psoriatic skin. Epidermal and dermal regions are annotated, with most keratinocytes expected in the epidermal region.

Although BLASE can deconvolute the 10x Visium spots along pseudotime, it struggled to attribute spot mappings with high confidence. One explanation for this could be that the low number of transcripts in each spot introduce noise that BLASE cannot overcome. Alternatively, the mixture of cell types may be too heterogenous for our approach to produce confident results. We also applied BLASE to another dataset [3] which yielded no results, as the the spot size of the Visium platform was too large to provide a clear picture of the process (see Figure A.4). The spot size for the Visium platform is around 50*µm*, and the epidermis typically has a thickness of around 100*µm* [22], giving only 2 spots of resolution to investigate a complex and dynamic process. Although BLASE does not generate results with convincing confidence for this case, we can see a clear trend of granular keratinocytes being predicted at the outermost layer of the epidermis, and less mature cells just below the outermost layer, reflecting the biology (see Figure A.4). Despite reflecting the general biology, BLASE seems to call spots in the upper half of the image as granular keratinocytes even though they are too deep into the tissue for this to be likely to be correct. It can also be seen that BLASE tries to calculate mappings for spots deep into the dermis, which is unlikely to have a high proportion of keratinocytes, which is likely an artefact of the method that was used to identify spots to deconvolute (by using high expression of keratinocyte marker genes in spots, described in detail in Methods). The phenomenon known as “spot swapping” where RNA does not stay localised to the spot it was originally within) could explain both the unlikely mappings, as well as the high expression of marker genes found deeper into the tissue than expected [23]. As ST technologies improve (for example 10x Visium HD), and other tissues are captured, BLASE will be able to map RNA-seq data to processes in spatial data.

### 3.4 Use Case 2: Identifying and labelling lifecycle stages

The lifecycle of the malaria parasite is complex, including life stages in both vector and host species. Part of this complex cycle is the red blood (or intraerythrocytic) stage, where the parasite infects red blood cells, multiplies and then bursts out of them to begin the cycle anew, exponentially increasing the population in the host [24]. In the human parasite, *P. falciparum*, this stage of the lifecycle repeats every 48 hours and is of special interest as it causes the worst pathology of malaria. These stages are: ring, trophozoite and schizont. There are several microarray [25, 26, 27] and bulk RNA-seq datasets of this cycle [28, 29]. These historic experiments with time-series microarray and bulk RNA-seq experiments have often revealed key insights into the transcriptional signatures underlying the process, even without the advantages of scRNA-seq. Several scRNA-seq datasets also exist [30, 2] and have provided deeper insights into *Plasmodium* biology.

Here, we will evaluate the mapping of a given bulk sample to its position in the cycle, comparing different genes and bins in BLASE, as well as existing methods, to show that this information can be used to annotate scRNA-seq data using existing bulk RNA-seq data.

#### 3.4.1 Evaluation of mapping

This example demonstrates BLASE’s utility for mapping a process in single-cell data from better annotated bulk RNA-seq data. In the Dogga et al. 2024 dataset used here as a reference, a similar method of correlation with bulk samples from four historic time series experiments [28, 26, 29, 27] were used to inform cluster annotation, in a more heuristic fashion than presented by BLASE. To validate BLASE on this dataset, we used microarray data of synchronised *P. falciparum* parasites sampled hourly over the 48 hours of the parasite’s intraerythrocytic lifecycle [26] and the scRNA-seq data of *P. falciparum* (a subset of 6000 cells) data from the Malaria Cell Atlas [2]. The optimal result would be that each bin of the pseudotime has a roughly even number of bulk samples mapped to it, moving across the cycle. Of note is that scRNA-seq of Dogga may not include high coverage of ring stage parasites [31].

BLASE correctly assigned every bulk sample tested when using every gene, according to identifications given by Painter in their paper analysing the data (see Table A.2). In Figure 4 (a), we show the UMAP projections of cell types (as annotated in the Malaria Cell Atlas) and pseudotime bins (b), as well as the correlation heatmap of the results calculated by BLASE (c). This heatmap also includes annotations of expected cell types for the bulk samples and for the pseudotime bins based on the annotations from their original analyses.

**Figure 4.**
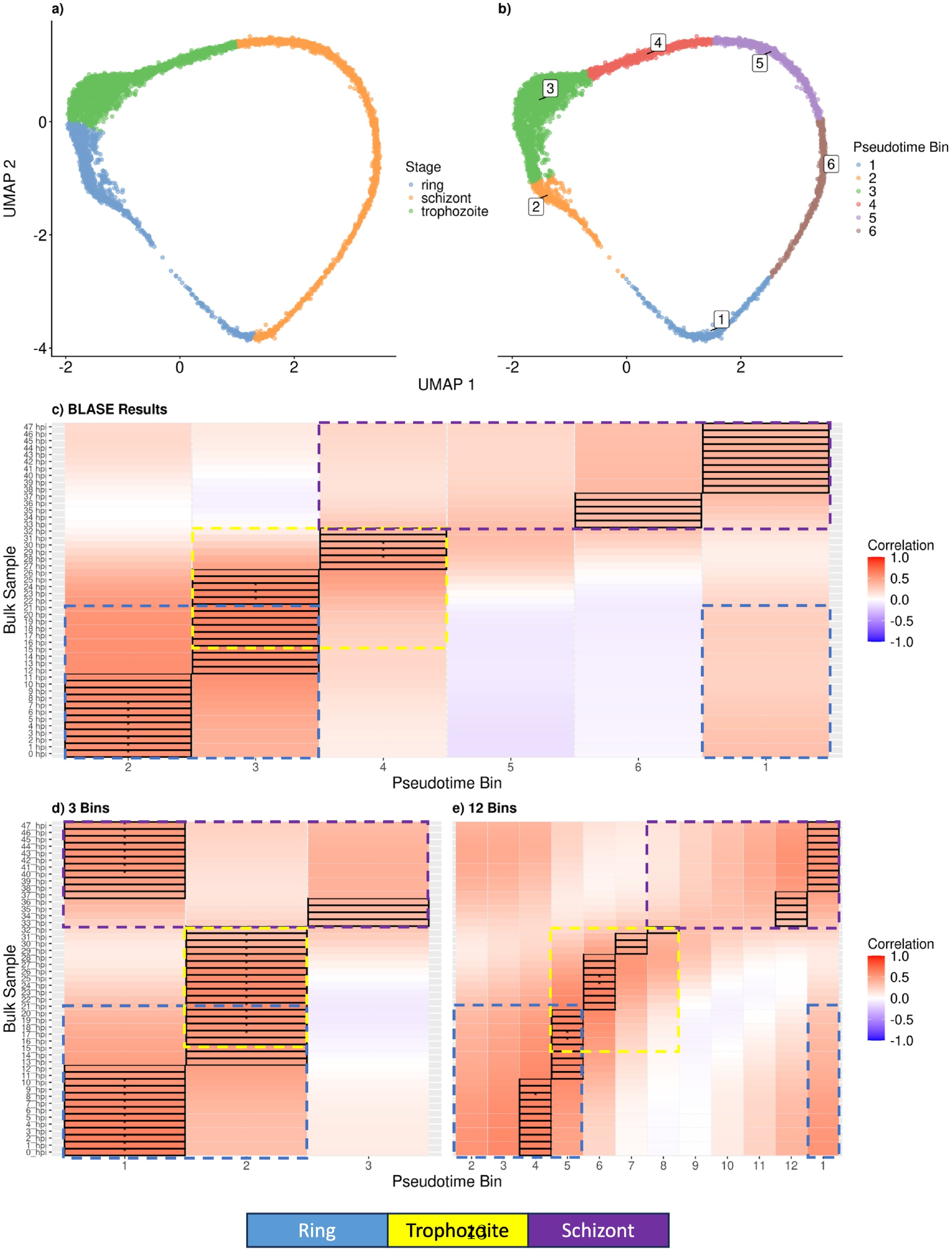
BLASE infers pseudotime on scRNA-seq reference trajectories using real single-cell and bulk datasets of *P. falciparum* intraerythrocytic life stages. a, UMAP embedded plot of the single-cell reference dataset, coloured by cell types annotated in the original analysis and b, pseudotime bins as assigned by BLASE. c, A BLASE results heatmap for this pair of datasets using 6 bins. See Table A.2 for the canonical annotations in detail. Pseudotime bins are shown starting at bin 2 to emphasise the progression predicted by BLASE. d,e, The same as c, but using 3 and 12 bins respectively.

The number of bins used by BLASE introduces an important trade-off between granularity and confidence. A greater number of bins increases the resolution of the results, however, this results in each bin becoming more similar to its neighbours. This increases the likelihood a sample will be incorrectly mapped to an adjacent (or further) pseudotime bin. BLASE is furthermore less likely to calculate that these mappings are confident due to the increased similarity. Reducing the number of bins will generally provide more confident results, but can be of reduced utility due to this loss of granularity. We show a comparison of this data mapped with varying numbers of bins in Figure 4 d, e. Further examples are given in Figure A.5.

Using 6 bins, we found that late schizont stage pseudotime bulks mapped to bin 1, reflecting that bin 1 is annotated in Dogga et al. as being a mix of schizont and ring stages. Early ring time-points mapped to bin 2, followed by late ring and early trophozoite stages in bin 3. Later trophozoite stages mapped to bin 4. No bulk samples mapped to bin 5: pseudotime is not necessarily linearly related to real time, and the parasites may have developed quickly over this part of the trajectory, before finally showing an early-mid schizont transcriptome in bin 6. This effect is exacerbated in the example where 12 bins were used for the mapping (Figure 4, e). In an ideal case, where pseudotime was linearly related to the passage of real time, we would expect to see an even number of hourly time-points in each bin, sequentially moving through the pseudotime. However, as can be seen this is not the case, and is an important caveat to bear in mind when making use of both BLASE, and more widely, trajectory inference methods in general. It is also important to note that this effect could be caused by the single-cell reference having a low number of cells representing these time-points, introducing bin-specific noise which results in mappings to nearby time-points.

#### 3.4.2 Gene selection

We additionally applied existing cell-type deconvolution methods and BLASE with varying gene lists. Four lists of genes were used for the BLASE comparison:

- All genes: This gene list allows BLASE to use every gene in the data.
- TradeSeq: 800 genes selected by using TradeSeq to find genes which are differentially expressed over the trajectory.
- Gene Peakedness: 800 genes found by using BLASE’s gene selection method to find the most highly peaked genes in the dataset.
- Gene Peakedness Spread: 919 genes selected as with Gene Peakedness above, but using the `gene_-_peakedness_spread_selection` function to select the most peaked genes over multiple slices of pseudotime, to ensure that the genes represent the whole trajectory.

All “confident” calls made by BLASE were correct, and additionally, all non-confident predictions were correct. The TradeSeq gene list had the highest number of confident mappings (30), suggesting that it is a good method for reducing the noise of genes which are not useful for determining the pseudotime of a cell. Using every gene gave 27 confident mappings, whereas both gene peakedness methods found fewer confident mappings (15 without spread, 21 with). BLASE outperformed all the other methods except for DWLS which equalled BLASE by also correctly mapping all 48 samples. CIBERSORTx performed well, only incorrectly mapping 1 sample. MUSIC and MeDuSA both performed poorly, with only 34 and 33 correct mappings respectively (Table 1, Table A.2, Figure A.6).

**Table 1:** Comparison of BLASE using four different gene selection methods (TradeSeq, All genes, Gene Peakedness and Gene Peakedness Spread), and four other deconvolution tools (CIBERSORTx, DWLS, MeDuSA, MuSiC) when trying to identify the pseudotime bins of the Painter et al. 2018 dataset on the single-cell reference (Dogga et al. 2023). We calculate the correct calls using the mappings in Table A.2. For BLASE mappings, the number of “confident” calls is given, and how many were correct.

| Method | Correctly Confident | Correctly Predicted |
| --- | --- | --- |
| **BLASE (1300 genes, TradeSeq Selection)** | **30/30** | **48/48** |
| BLASE (All Genes) | 27/27 | 48/48 |
| BLASE (1300 genes, Gene Peakedness Spread Selection) | 21/21 | 48/48 |
| BLASE (1300 genes, Gene Peakedness Selection) | 15/15 | 48/48 |
| DWLS (By majority call) | NA | 48/48 |
| CIBERSORTx (By majority call) | NA | 47/48 |
| MUSIC (By majority call) | NA | 34/48 |
| MeDuSA (By majority call) | NA | 33/48 |

This example demonstrates that BLASE not only outperforms existing cell-type deconvolution tools at this task, but that BLASE can do so with perfect accuracy. BLASE, even with less effective gene lists, substantially outperformed some of the other tools.

#### 3.4.3 Cell annotation

Similar to the Dogga et al. 2024 dataset used here, which made use of correlation with bulk data to annotate their cells, we have implemented in BLASE the option to annotate the cells taken from mapping bulk RNA-seq samples. In Figure A.7 we show how we mapped the 48 hour Painter time-series data and two other datasets to the Malaria Cell Atlas *P. falciparum* dataset.

In this example, the perfect scenario would be to have a 48-bin scRNA-seq reference whose pseudotime bins were split exactly according to real time, that is, we would expect the cells in pseudotime bin 1 to be only cells from the first hour of growth, and so on, with each pseudotime bin representing one hour of growth. However, the trajectory of this scRNA-seq reference displays compressions in pseudotime, where in some regions, an increase of pseudotime is not directly proportional to the real time which has elapsed. Figure A.7 shows this in c, d and e, where each there is not a one-to-one mapping from pseudotime bin to bulk sample, so some bulk samples are the best match for multiple pseudotime bins, and some bulk samples are completely skipped. Furthermore, BLASE can include the correlation of each of these, giving a confidence value to each “annotation.” Since BLASE is implemented as a tool, it is extremely convenient to repeat this analysis with two other bulk datasets [29, 32], which can help identify areas of uncertainty which may be due additional attention when using BLASE for annotations.

### 3.5 Use Case 3: Detecting developmental shift in knock-out and correcting differential expression

In many knock-out and knock-down cell lines development can be impacted or even arrested at certain points. This can make RNA-seq experiments extremely challenging, as comparisons with wild-type cells can give results that are heavily biased simply by differences in cell development, obscuring other changes induced by the mutation (Figure 5 a).

**Figure 5.**
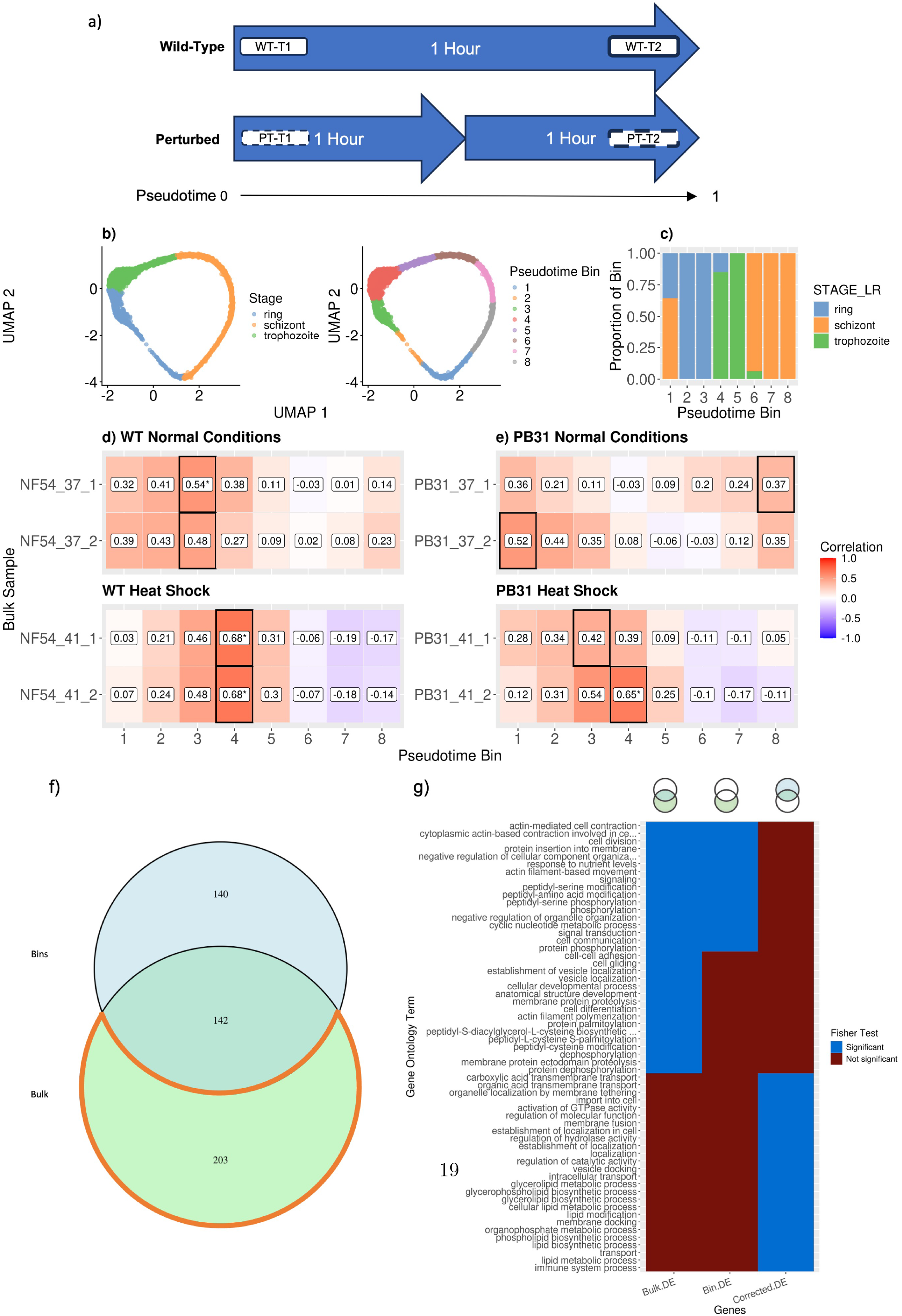
BLASE can reveal new insights from published heat shock data. a, A schematic showing the way in which a bulk RNA-seq timecourse experiment with two conditions can be confounded by the effect of applying another condition. Line one is wild-type, and line two is subjected to a perturbing condition. The perturbed population is developmentally delayed, taking two hours to progress to the transcriptional state that the wild-type reaches after one hour. b, UMAP embedded plots of the single-cell reference dataset, coloured by cell type and pseudotime bin. c, The proportions of each cell type found in each pseudotime bin. d, A BLASE results heatmap of wild-type *P. falciparum* following exposure to heat shock (41^°^C) or kept at 37^°^C, cultured over three days. e, The same as d, but for a *P. falciparum* isolate with the *LRR5* gene knocked out. Note that in both d and e, the sample subjected to heat shock is predicted to have developmental differences with the wild-type. f, A Venn diagram showing the numbers of genes found to be DE (between PB31 under normal and heat shock growth conditions) in the comparison of the reference bins (Bins), between the bulk samples (Bulk), and the intersection of the two. g, GO enrichment analysis correction with BLASE for the comparison of PB31 under normal or heat shock growth conditions. A heatmap of the GO terms found from each list, which were found to be significantly enriched exclusively in the GO term enrichment from the corrected or uncorrected list of genes.

During its different lifecycle stages, the malaria parasite undergoes extreme and fluctuating thermal conditions - most importantly the high temperatures during malarial fever - and has evolved mechanisms to survive and thrive in such a hostile environment. Understanding the heat-shock response might yield novel treatment targets: Zhang et al. [33] found through their QIseq approach [34] a mutant that is sensitive to heat-shock (labelled PB31) due to a Piggybac insert in the *LRR5* gene (PF3D7 1432400). This gene contains leucine rich repeats which suggest that it may be involved in protein-protein interactions [35, 36]. The authors hypothesised that it would grow differently under heat shock conditions compared to the wild-type. To investigate, they performed RNA-seq of both wild-type and mutant isolates after 3 days of culturing, either with heat-shock regularly applied (simulating the conditions of malarial fever), or without. In order to understand the true impacts of these conditions, it is important to have confidence that the samples are comparable with one another.

#### 3.5.1 Detecting growth differences

We hypothesised that knocking out this gene could affect the growth rate of the parasites, confounding the analysis of bulk RNA-seq samples taken at the same time point. We used the Malaria Cell Atlas as a reference [2] (Figure 5, b, c).

BLASE identifies a clear change in development between normal culture conditions (bin 3) and heat shock in the NF54 reference isolate (mapped to bin 4) (Figure 5, d). The PB31 mutant under normal conditions also has a developmental difference with the reference under normal conditions, as the two biological replicates map to bins 1 and 8. As the *P. falciparum* lifecycle is 48 hours, this difference with the WT isolate (NF54) is as large as 18 hours. Under heat-shock the PB31 isolate appears to be at a similar time point to the NF54 under the same condition (Figure 5, e). Although other methods exist to detect the timing of malaria parasites [37], BLASE has the advantage that bins can be adapted according to requirements and can use a continuous pseudotime reference. Further, it can be noted that overall the WT parasites are more synchronised, as the biological replicates mapped to the same bin and the mapping scores are higher at the peak-call. However, in the heat-shock conditions, the parasites are more synchronised as the mapping score is higher and less widely distributed across neighbouring bins. This is to be expected, as heat-shock can be used to synchronise intraerythrocytic *P. falciparum* [38].

These differences could mean that these samples are not as comparable as they outwardly appear. Normal developmental differences may be the strongest signal when performing comparisons such as differential gene expression analysis, instead of the underlying transcriptomic process of heat shock response. By using BLASE, these discrepancies can be identified and subsequently accounted for, aiding researchers in understanding the most meaningful differences.

#### 3.5.2 Correction of effect of growth in DEG

Due to the different growth phenotypes in the isolates, a normal differential expression (DE) analysis will primarily return cell-cycle genes, rather than differentially expressed genes (DEG) related to the phenotype. To showcase this, we performed a differential expression analysis of pseudotime bin 1 against bin 4, detecting 581 genes upregulated in bin 1, which then informed GO term enrichment analysis, yielding three top terms of: “symbiont entry into host”, “cell motility” and “biological process involved in interspecies interaction between organisms”, which are attributed to life-stage changes (Additional file 1). These signals hamper efforts to identify real biological differences, and we propose that BLASE can be used to correct for that error by ignoring the DEG found from cell cycle development differences, by subtracting them from samples taken at different time-points.

This approach is needed as we want to compare the differences between the normal PB31 condition and under heat-shock. To understand the phenotype differences due to the heat-shock treatment, we must correct the DE analysis of genes due to the impact of the cell cycle genes. As shown before (Figure 5, e), parasites of the PB31 isolate under normal conditions map to bins 1 and 8 and those cultured with heat-shock to bins 3 and 4. Performing DE analysis of those two conditions in the scRNA-Seq gives 282 genes exclusively due to cell cycle differences (Figure 5, f, Additional file 2). This number is lower than in the analysis of bin 1 against bin 4 above, as here we use a bulk DE approach to mirror the bulk comparison, which returns fewer false positives. Performing the DE analysis between the normal and heat-shock condition of PB31 returns 345 genes, (Additional file 2).

Overall, there is an overlap of 142 genes between these two comparisons (Figure 5, f), leaving 203 genes only in the bulk DE analysis following correction. To explore the function of the genes in the bulk, single-cell pseudotime bin (bin), and post-correction (corrected) DEG lists, we performed GO enrichment analysis of each list. In figure (Figure 5, g) we can see three different patterns. First, GO terms like “Cell division” or “signalling” which are attributed to the cell cycle, as they occur in both the bulk and bin DEG comparison. However, some GO terms, “cell adhesion” or “cell differentiation” are found in the enriched GO terms of the bulk genes, but not the bin or the corrected, although they are cell cycle differences. The reason for this is that due to the correction, some GO terms are no longer significant. Finally, we can see GO terms just in the corrected process, such as “transmembrane transport”, and “lipid metabolism” which are informative for the impact of the heat-shock on the mutant (Additional file 3). Here, BLASE reveals previously obscured information from the bulk RNA-seq data, highlighting biological insight.

In summary, here we have shown two more use cases for BLASE. First, BLASE confidently maps bulk transcriptomics data to the malaria intraerythrocytic cycle, and secondly how scRNA-seq data and pseudotime can be used to correct differential expression analysis in asynchronous samples to detect hidden biology.

Although there are other methods in the malaria literature that propose linear models to correct the data [39], we note that they also rely on reference data which are generally discrete, and they rely on the fact that plasmodium blood stages are cyclic, with gene expression tightly regulated along this cycle. Other life stages of this parasite, other pathogens and most other cells in general, are not. Therefore, compared to existing methods from the malaria, we propose a more general solution, which is both more robust and more flexible.

### 3.6 Runtime and implementation

We compared the running time of BLASE with the other tools (Figure 6, a). This showed that MUSIC was consistently the fastest method. CIBERSORTx and DWLS were the slowest in both cases. The computation done by BLASE scales by: *bins* × *samples* × *bootstrap iterations*. In the haematopoiesis example, with 5573 cells, 19178 genes and 5 samples, BLASE finishes within a few seconds, DWLS takes over 25 minutes. For situations where many bulk samples must be mapped, parallel execution can be used in BLASE to increase the speed of computation, such as in the case of our *P. falciparum* lifecycle mappings, which included 48 samples, with 5275 genes in 6000 cells. The largest component of BLASE’s running time is the bootstrapping step to calculate confidence, with a default of 200 iterations. However, even when single-threaded, BLASE completes running quickly. Without being as capable of scaling up in this way, using BLASE to analyse over 1000 Visium spots would take a prohibitive amount of time.

**Figure 6.**
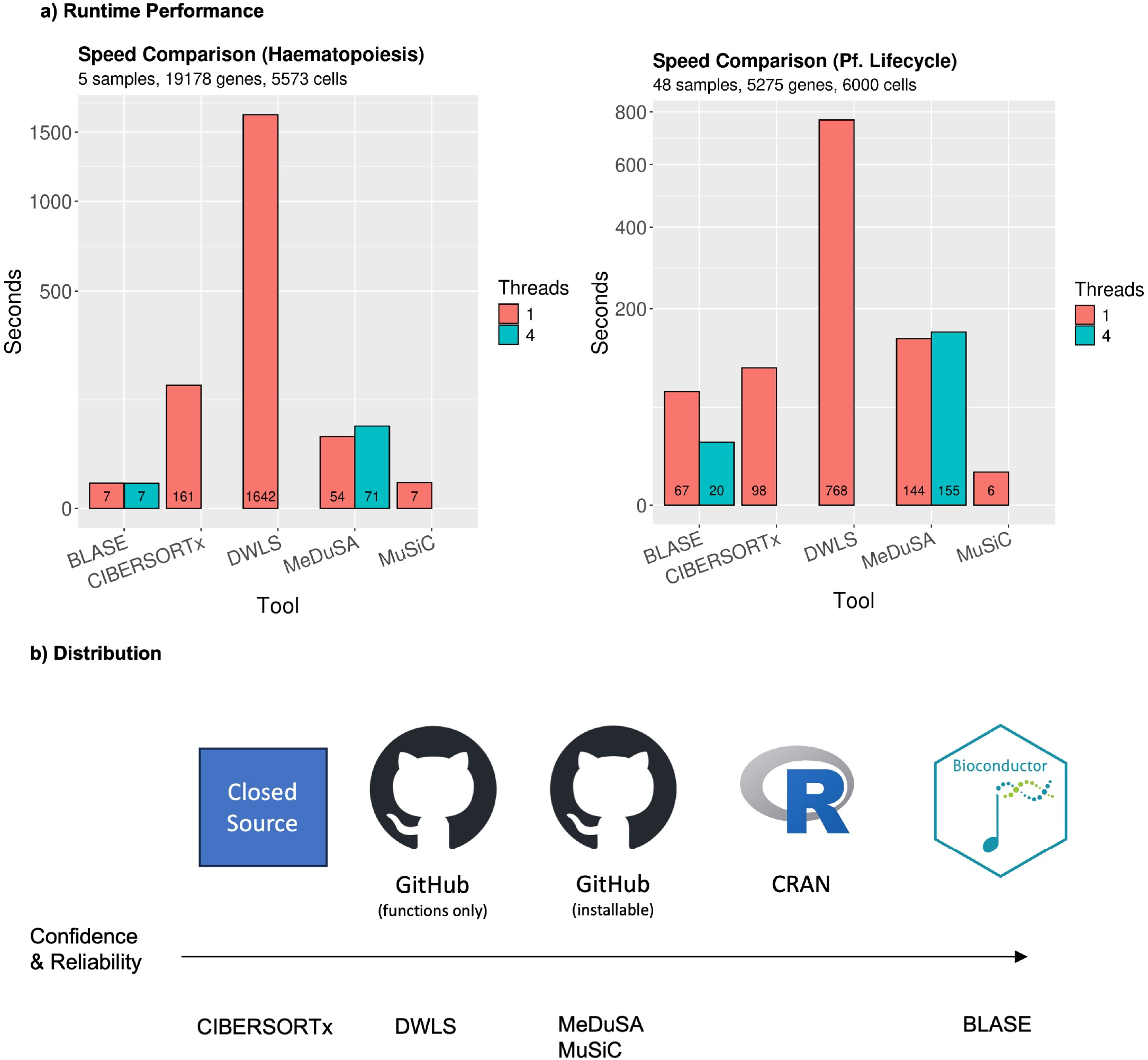
a, Runtime of BLASE and other tools in seconds for two example datasets, the first a haematopoietic myeloid lineage, and the other the asexual differentiation of *P. falciparum*. b, The distribution methods employed by BLASE and other tools, including closed-source, open-source, and packages installable through GitHub, CRAN, or Bioconductor. Packages available on CRAN and Bioconductor must meet good-practice requirements, as assessed by each organisation. R logo used under license (https://www.r-project.org/logo), Bioconductor license used with permission.

BLASE will be available for installation via Bioconductor, which guarantees that the code matches Bioconductor’s requirements for quality (https://contributions.bioconductor.org) – including version control (through GIT), complete documentation (built with pkgdown [40] and distributed through GitHub Pages), following R coding practices and in excess of these requirements, unit testing (testthat [41]). The source code is freely available, and is released under a GPLv3 license (Availability of Data and Materials). In Figure 6 b, we show how other similar tools have been released. CIBERSORTx is released as a closed-source website and docker container which requires a token to run. DWLS is released as R scripts which can be run locally. MeDuSA and MuSiC are both installable from GitHub, but are not released through a package management system such as CRAN or Bioconductor, and as such are not guaranteed to meet the respective standards of those organisations.

## 4 Discussion

Although the deconvolution of bulk RNA-seq through scRNA-seq per se is not novel [5, 42], none of the other tools allow the user to perform all of the above use cases out of the box. Further, BLASE generally matches or outperforms those existing tools, in both simulated and real pseudobulked datasets, where we have a ground truth to validate against. As many deconvolution tools focus on cell-type proportion deconvolution, we cannot provide a truly fair comparison. However, Medusa is more comparable to BLASE, as it also assumes pseudotime is available to map to, and we show that BLASE outperforms MeDuSA. Again, the cell type deconvolution tools generate statistics for cell-types proportion, and we generate statistics for the best call.

We suggest several methods for the selection of temporal marker genes: using every gene available, adapting existing methods such as TradeSeq, or using BLASE’s gene peakedness metric. By remaining agnostic to the set of genes being used, we invite researchers to make use of their existing expertise in the biological processes they study, instead of relying on purely automated techniques. BLASE provides a statistical framework for evaluating confidence in its results through bootstrapping. Not only does this allow BLASE to predict confident mappings, researchers can also look into the confidence intervals for each mapping.

The applications of BLASE are not limited to bulk transcriptomics. With the advent of ST and methods such as GeoMX and 10x Visium, the field will generate “spatial bulk” data from tissues. We have shown that with the right skin dataset, we can see development in the epidermis mapped onto the spatial information these datasets provide. Although the experiments did not yield novel insight into psoriasis, we have shown that BLASE has a multitude of potential applications.

We applied BLASE to different published datasets to showcase these applications. BLASE accurately identified the correct cell-type mappings for a malaria dataset, showing that it is possible to use bulk RNA-seq data to annotate single-cell data. For the latter task, we can also annotate the scRNA-seq bin better than in the MCA examples (Figure A.7), however, there is still room for improvement. It should be noted that the reference datasets might not have all the life-stages perfectly represented, and BLASE, as well as the other cell-type deconvolution methods evaluated here rely on the reference dataset being correct and appropriate. With more atlas-size datasets becoming available, the BLASE approach will become more robust.

In the second malaria example, using Zhang et al. 2021 [33], we show how BLASE can reduce the impact of developmental genes associated with a trajectory, so that we can observe the impact of the heat-shock in the mutant. Although GO term enrichment analysis is not a perfect tool, especially in pathogens with many genes of unknown function, we can see that by filtering the input datasets, more relevant GO-terms associated to heat-shock are found. First, it might be unintuitive that those GO terms were not found in the comparison of the bulk, as the genes of interest are still there. However, when giving GO a large list of genes, the number of multiple tests increases, making some relevant GO-terms disappear. Therefore using our approach can highlight novel biology.

Finally, we would like to emphasise the modern and robust implementation of BLASE. We believe that software distributed for use in academia, like industry, must be robust and reliable. A variety of methods can be used to ensure that software released meets the needs of those using it, and different levels of effort should be applied depending on how widely the tool will be distributed [43].

## 5 Conclusion

Integration of different modalities in omics is always challenging. When performing transcriptomic experiments, single-cell sequencing offers many opportunities, however, it is more technically challenging and is many times more expensive than bulk RNA-seq. Here we introduce BLASE which enables the integration of single-cell and bulk RNA-seq data, by leveraging the fact that in biology many datasets have an underlying trajectory, such as the cell cycle, differentiation or growth. We propose three different use cases. First, the determination of the pseudotime of bulk samples against a single-cell reference. Second, BLASE can also be used to accurately annotate scRNA-seq data using existing annotated bulk RNA-seq, and finally, we can use the mappings of bulk data onto a single-cell reference to correct DE analysis error caused by asynchronous samples.

In conclusion, we present a versatile tool for the community to enable the re-analysis of existing transcriptomics data in the context of pseudotime, leading to novel insights without the investment of time and money in new scRNA-seq experiments.

## 6 Methods

Here we present details of how BLASE discretises pseudotime and maps bulk samples onto scRNA-seq trajectories using a best correlation approach. A broad description of the BLASE algorithm is given in Figure 1, and Figure A.9 is a detailed, step-by-step flowchart of the algorithm. BLASE was programmed in R, and will be installable through Bioconductor. Further detail can be found in the code, maintained on GitHub (Availability of Data and Materials).

BLASE provides methods for splitting a single-cell RNA-seq (scRNA-seq) reference into pseudotime bins, evaluating hyperparameters, mapping a bulk RNA-seq sample to the best matching pseudotime bin, and then plotting these results. The algorithm for mapping a bulk RNA-seq sample to a pseudotime bin is given in subsection 6.1.

The typical workflow (Figure 1), starts with a scRNA-seq dataset, where each cell has been assigned a “pseudotime” value designating its relative progress through a biological process. A list of genes which are informative regarding the progression of a cell through the pseudotime linage is also required. This may be all available genes, or a subset, depending on the process of interest. BLASE provides methods for assisting in the determination of which and how many genes should be used (see subsection 6.2). A list of genes ordered by how informative they are about the pseudotime of a cell in the lineage is also required, methods for which are also provided by BLASE (see subsection 6.3).

BLASE takes a matrix of normalised scRNA-seq data, *X*, with *p* rows representing the genes, and *c* columns representing the cells. The cells are ordered according to their pseudotime, labelled *t*_*j*_ ∈ [0, *τ*], ∀*j* ∈ {1, …, *c*}, where *τ* is the maximum pseudotime. Normalised bulk data, **y**, of length *p* must also be supplied, with both *X* and **y** using the same set of *p* genes, 𝒢.

This method also makes use of hyperparameters. As the full set of genes does not necessarily need to be used in BLASE, calculations are performed on a subset of *p*^′^ genes, 𝒢^′^ ⊆ 𝒢. Also, the set of cells needs to be partitioned as 

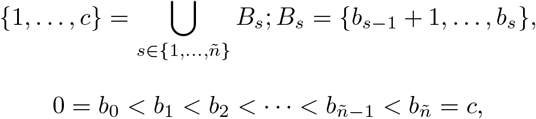

where ñ is the number of pseudotime bins. Methods to suggest 𝒢^′^, ñ and the *b*_*s*_ terms will be described in subsection 6.2, subsection 6.3 and subsection 6.4.

### 6.1 BLASE: Mapping

In order to identify the best match for a bulk sample within a list of pseudotime bins in the scRNA-seq reference dataset, BLASE scores similarity with Spearman’s Rho, and calculates confidence intervals using bootstrapping.

Mapping is performed by the `map_best_bin` or `map_all_est_bins` functions for a single or many bulk samples respectively. The algorithm proceeds as follows for each RNA-seq sample (see Figure A.9 for a flowchart of the algorithm).

Because we are using 𝒢 ^′^, we may not be using the full scRNA-seq data, *X*, but rather a matrix produced by keeping only the genes in 𝒢 ^′^, which we label *X*^′^. Similarly, the bulk data, **y**, is transformed to produce **y**^′^. Having binned the cells according to pseudotime, we produce pseudotime bulk vectors for each bin by adding the normalised RNA counts across the bin for each gene. We define this by 

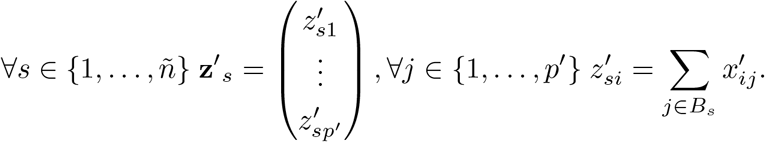

This is used to calculate the correlations between **y**^′^ and 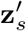 for all bins *s*, and the confidence intervals for these correlations.

Bootstrapping is used to calculate the confidence intervals. For all *s* ∈ {1, …, ñ} and *l* ∈ {1, …, *k*}, where *k* is the number of bootstrap samples, a sample of size *p*^′^ is drawn from 𝒢^′^ with replacement, labelled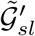. By subsetting (with duplicates for repeated genes) 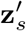 and **y**^′^ with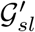, we generate the vectors 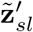 and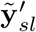. The correlation between 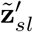 and 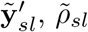 is then calculated.

By iterating over all *l*, a set 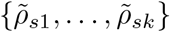 is calculated. The confidence interval is the 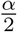 and 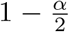 quantile of this set, labelled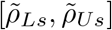. The point estimate of the correlation between 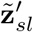 and 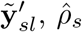 is obtained by calculating the Spearman correlation between the non-bootstrap 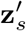 and **y**^′^ as usual.

We define the most suitable bin as that with the highest correlation with the bulk data, while the bootstrap intervals are used to establish whether we are confident in this assessment. The best and second best bins are defined as ŝ_1_ and ŝ_2_ respectively, where: 

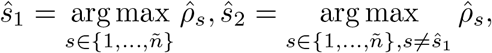

and we say that a bin is confidently mapped by *ŝ*_1_ when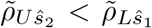. This means that a mapping is considered confident when the pseudotime bin with the highest correlation to the bulk’s lower bound is still greater than the upper bound of the next best mapping pseudotime bin.

The implementation of bootstrapping uses the `spearman.ci` function from the RVAideMemoire package [44].

### 6.2 BLASE: Hyperparameter selection

In order to help with making informed decisions about trade-offs between the number of bins and genes used for mapping, BLASE provides functions that calculate heuristic measures of how to select these cutoffs. These make use of a measure of how particularly a bin is associated with pseudotime, which we call convexity. Subsequently, the number of bins and the list of genes are used in the mapping step.

The `find_best_params` function takes a range of possible gene and bin counts, and calculates the minimum convexity, mean convexity, and number of confident mappings (subsection 6.1) for each combination. It should be noted that this can only take into account the reference dataset, and may not be an accurate representation of the best values for a given bulk sample.

𝒢 = {*g*_1_, …, *g*_*p*_} is assumed to be in descending order of “goodness” (by the methods described in subsection 6.3, or by another method deemed suitable by the researcher using BLASE). The objective is to calculate an optimal number of genes, *p*^′^, and bins, ñ.

For each pair of hyperparameters considered, 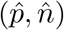, the reference data is pseudobulked into 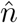 bins. With a portion of the reference data pooled into 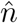 pseudobulk data, mapping on each pseudobulk bin can be performed as described above using the first 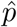 genes,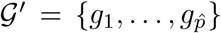. For each pseudobulk sample 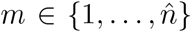,we calculate its convexity, 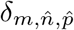, which is a measure of association between a gene and a bin. This is defined by 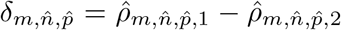,where 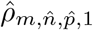is the highest correlation calculated between bin *m* and the pseudobulk data, and 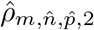nis the second highest, for bin and gene counts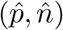.

The suitability of any 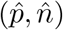 can now be assessed by the mean convexity over bins, 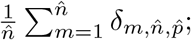 the minimum convexity, 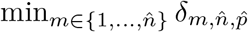; and the proportion of confident bin assignments.

The `gene_selection_matrix` function plots log normalised expression of genes in cells. The y-axis represents genes, ordered by the pseudotime of their peak expression, the x-axis represents cells, ordered by pseudotime. This can be used to assess how well differentiated pseudotime is by a selected list of genes.

### 6.3 BLASE: Gene selection

BLASE provides several functions to allow researchers to make informed decisions about selecting which genes should be utilised by BLASE. However, use of these methods is optional and they are provided only as a convenience and recommendation. Researchers will understand their research area best and can decide if these methods are appropriate. A comparison of these methods across datasets is shown in Figure A.10

#### 6.3.1 Gene peakedness selection

Genes which have a peak of expression at certain points in the pseudotime trajectory are useful as indicators of the progress of a cell through the process. When a cell has a high expression of this gene, it is likely that the cell is at that given point in pseudotime. These genes can be found by attempting to identify how “peaked” the expression of the gene over pseudotime is within the reference dataset. First a GLM is fitted to the normalised expression over pseudotime in order to calculate a smoothed expression over pseudotime. Then, within a given window around the peak of the smoothed expression, calculate the mean normalised expression within and without the window. A score is calculated as the mean normalised expression of the gene within the window divided by the mean expression outside the window. A higher score is considered more peaked and therefore more useful.

The normalised expression data for each gene *i* ∈ 𝒢 is smoothed with respect to pseudotime using the R package mgcv [45]. This fits a Generalised Linear Model (GLM) with a log link function estimating the mean normalised expression using a cubic spline on pseudotime (with a default of 10 knots). This gives a prediction function, 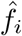, which maps from pseudotime to a normalised expression value. We calculate the normalised expression values on a regular grid over pseudotime, defining 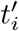 as the point at which the expression value is maximal. This estimates the pseudotime at which expression peaks. In order to split cells into inside-peak and outside-peak, the width of the peak as *w*, where by default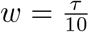. The cells inside the peak are 

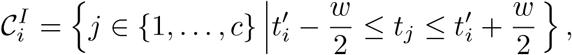

with all other cells in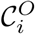. The inside-peak and outside-peak means are defined as: 

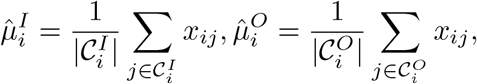

and the ratio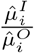 is used as a measure of gene peakedness, and it is recommended that the gene list is arranged in descending order of this ratio.

#### 6.3.2 Gene peakedness spread selection

To account for the possibility that the genes with a high peakedness ratio are concentrated at certain points in the pseudotime, genes can be selected in a way that enforces selection of the genes with the highest ratio throughout the trajectory. A function is provided for this in BLASE: `gene_peakedness - spread_selection`, which implements the algorithm outlined in Algorithm 1.

##### Algorithm 1

Selecting genes with high peakedness from over entire pseudotime.

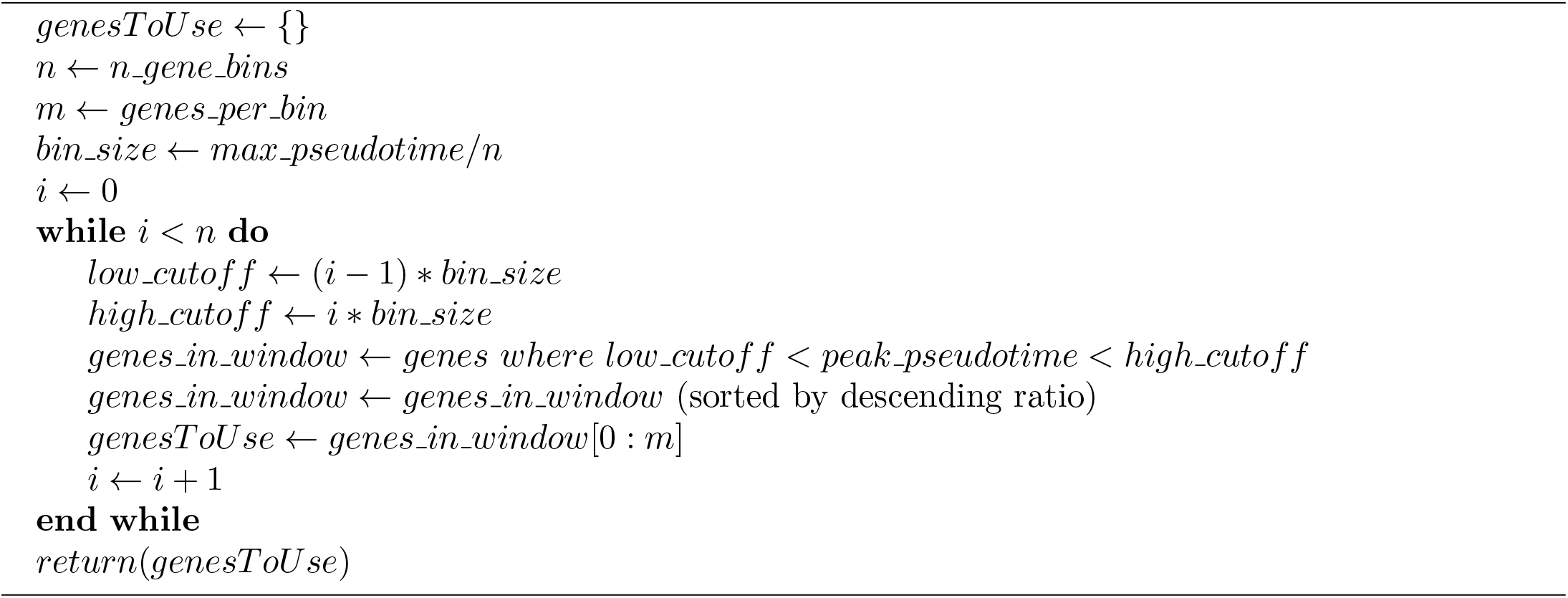

#### 6.3.3 TradeSeq

Another method for selecting genes which may be useful utilises TradeSeq’s `associationTest`, which finds the genes which change over pseudotime. The `associationTest` function can be tuned to identify genes which have different pseudotemporal patterns (using the contrastType parameter): differing at the start, end or between knots.

BLASE provides a method for selecting a number of genes with the largest differences across pseudo- time from TradeSeq’s association test, following the algorithm outlined in Algorithm 2, implemented in the `get_top_n_genes` function.

##### Algorithm 2

Selecting top genes from TradeSeq `associationTest` result

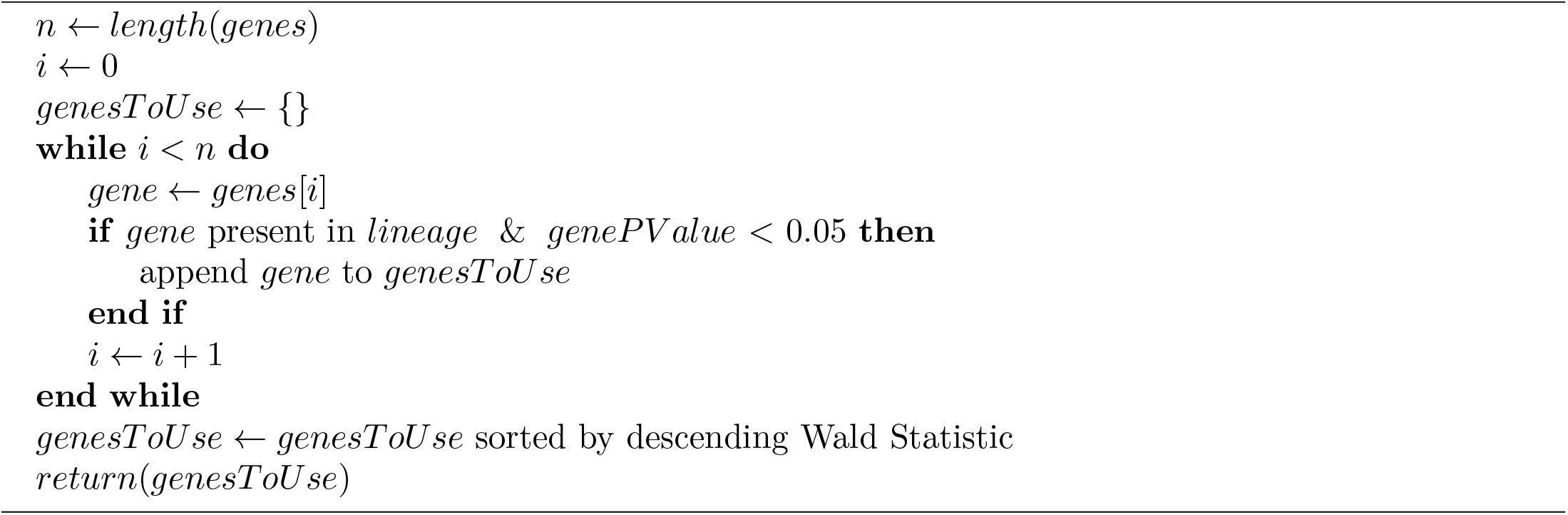

#### 6.3.4 All genes

In some cases, it may be preferable to use every gene available, and BLASE will accept any list of genes.

### 6.4 BLASE: Assigning pseudotime bins

BLASE provides two methods for the selection of bins (see subsection 6.2 for discussion of how to select the best number of pseudotime bins for a dataset). See Figure A.5, Figure A.11 for schematics of these methods. This occurs when a `BlaseData` object is created (typically using the `as.BlaseData` function), but can also be applied to an existing `SingleCellExperiment` object using the `assign pseudotime bins` function, enabling further analysis or visualisation.

#### 6.4.1 Pseudotime range bin assignment

Pseudotime range bin assignment assigns cells to bins on the basis of their progress through pseudotime. This method attempts to ensure that each bin covers a constant window of pseudotime, at the cost of variable numbers of cells per bin. Generally, this is the recommended method, however BLASE requires that every pseudotime bin contains cells, and there are some cases where bins will be created without containing cells when using this method.

Formally, in this method, all bins cover the same length of time in pseudotime, i.e.,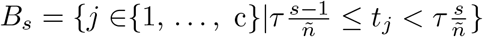.

#### 6.4.2 Number of cells bin assignment

Number of Cells bin assignment attempts to keep the number of cells per bin constant, at the cost of permitting varying pseudotime widths for each bin. This method can be vulnerable to bias from pseudotime abundance differences. For example, when one region of pseudotime has a much greater number of cells than another, it will produce proportionally more pseudotime bins. This means that the region with many cells will have many pseudotime bins, resulting in pseudotime bins in this region which may have very similar transcriptional signatures. This could affect confident calling in that region. In spite of these considerations, this method can be useful for datasets where certain pseudotime ranges have low or zero cell populations.

To split an even number of cells into each bin, the following is defined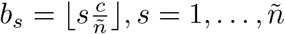.

### 6.5 Annotation of scRNA-seq using time-series bulk RNA-seq

In order to use bulk RNA-seq of known populations to annotate scRNA-seq cluster cell-types, BLASE implements `annotate sce`, which follows Algorithm 3.

### 6.6 Generation of scRNA-seq datasets

A copy of each of these objects, and code to reproduce them is provided on Zenodo (subsection 7.1).

#### 6.6.1 Generation of simulated dataset

To generate the in-silico simulated dataset for verifying BLASE can map pseudobulked data with a known ground-truth, Dyngen [17] was used to generate 2000 cells with 400 target genes, 50 transcription factors and 50 housekeeping genes, based on the “regulatorycircuits_04_mesenchymal mixed” feature net-work and “zenodo_1443566_real_silver_trophoblast-stem-cell-trophoblast-differentiation_mca” experiment counts, over a simulated time of 1000. “Cell type” clusters were identified using the Louvain algorithm in the bluster package [46].

#### 6.6.2 Adaptation of haematopoiesis scRNA-seq

To adapt the Mende et al. 2020 Haematopoiesis scRNA-seq dataset [18], the h5ad file was downloaded from the Human Cell Atlas [47]. It was then converted to a Seurat v5 object [48] by saving the counts matrix in CellRanger format, and the metadata as a tab separated value (tsv) file.

##### Algorithm 3

Annotating scRNA-seq cluster cell-types with BLASE results

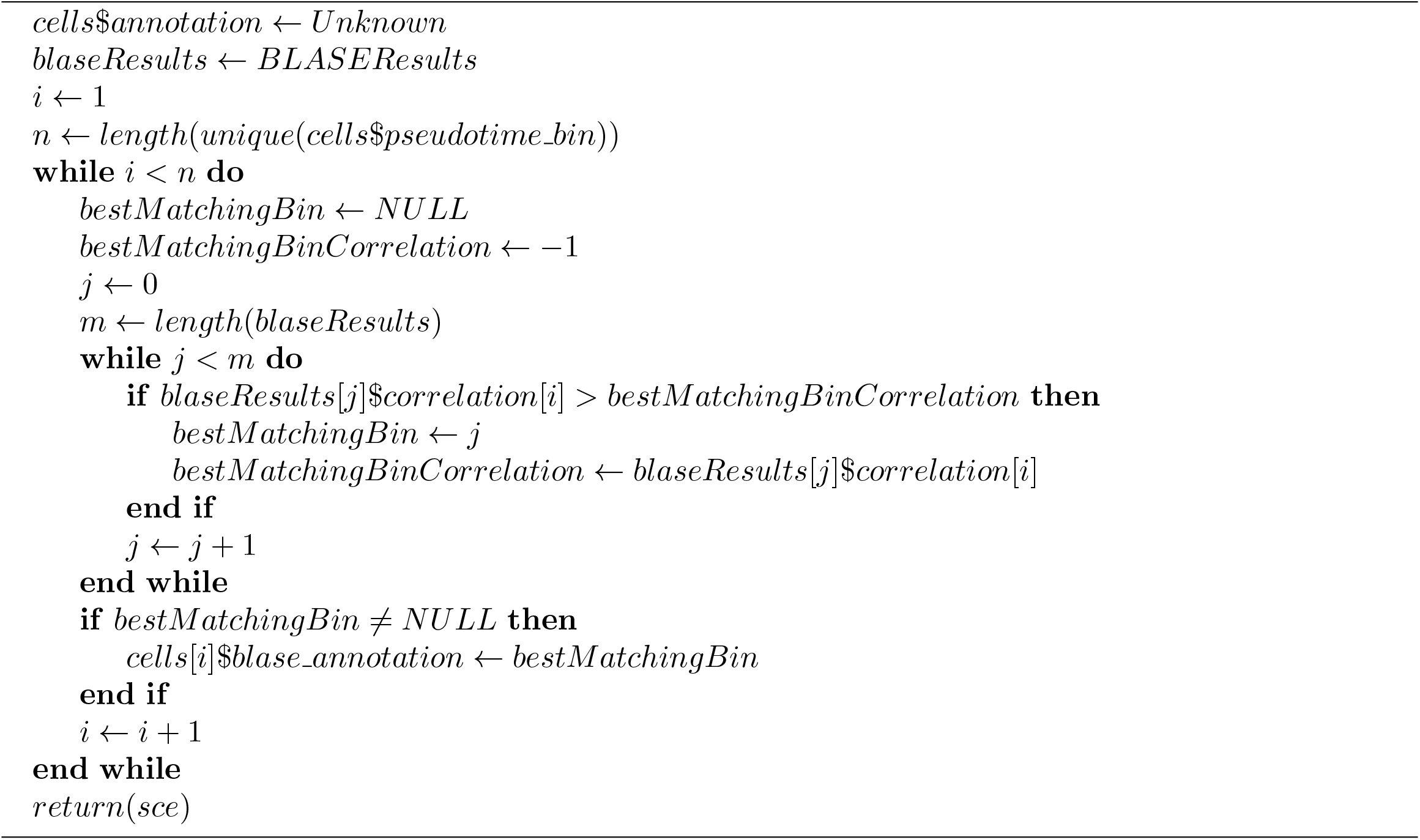

Quality control was performed for each sample with the below parameters:

| Library | Min. features per cell | Max features per cell | Max % Mitochondrial |
| --- | --- | --- | --- |
| SIGAD9 | 700 | 3500 | 5 |
| SIGAE9 | 700 | 3000 | 5 |
| SIGAF9 | 700 | 3000 | 5 |
| SIGAG9 | 700 | 3000 | 5.5 |
| SIGAH9 | 700 | 3200 | 5 |
| SIGAB10 | 700 | 4200 | 5.5 |
| SIGAC10 | 700 | 4000 | 4 |
| SIGAD10 | 700 | 4000 | 5 |
| SIGAA12 | 700 | 4000 | 5.5 |
| SIGAB12 | 700 | 4500 | 6 |
| SIGAD12 | 700 | 4000 | 5.5 |

Following Quality Control filtering, the data were normalized (`NormalizeData`), Variable Features were found (`FindVariableFeatures`, selection.method vst, nfeatures 2000), and scaled (`ScaleData`).

Principal component analysis (PCA) was then performed, and 15 principal components (PCs) were taken for further analysis. Integration was performed to merge each library in an unsupervised approach with STACAS [49], using the 15 PC embeddings and 1000 anchor features.

Nearest neighbours were found based on the first 15 PCs, and clusters found with a resolution of 0.1 (`FindNeighbors`, `FindClusters`). A PHATE embedding was generated for the integrated PCA embeddings using phateR [50]. 12000 cells were randomly selected from cells in the Stem Cell - Myeloid lineage, selected for using marker gene signatures:

| Cell Type | Markers |
| --- | --- |
| Stem Cell | SOCS2, MLLT3, HOPX |
| Myeloid Progenitors | ENO1, MPO, CEBPD |
| Myeloid | CD14, CD16, LYZ |

Cells along this lineage were reprocessed (as above) as a subset. Following this, a Single Cell Experiment object [51] was created, and Slingshot[10] was used to calculate the pseudotime from Stem Cell to Myeloid Cell. TradeSeq was run with 9 knots, selected using the `evaluateK` function.

#### 6.6.3 Adaptation of Malaria Cell Atlas *P. berghei* scRNA-seq

To adapt this dataset (from Howick et al. 2019 [30]), preprocessed data (pb-ch10x-set1.zip) was downloaded from the Malaria Cell Atlas (MCA) website (https://www.malariacellatlas.org/downloads/pb-ch10x-set1.zip). From the raw counts, log normalised counts were calculated using scran [52] (`computeSumFactors`, default parameters) and bluster [46] (`logNormCounts`, default parameters). The normalised counts were calculated as the exponential of the log normalised counts. A UMAP embedding was produced using scater [53] (`runUMAP`, default parameters), based on the principal components 1:2 provided with the object.

To calculate pseudotime, slingshot [10] was used (`slingshot`, “UMAP” reduced dimension, cluster labels based on MCA annotations, start cluster “ring”). TradeSeq [14] was run with 7 knots (selected by heuristic in the `evaluateK` function), and the `associationTest` results were calculated (lineages True, global False, contrast type consecutive).

#### 6.6.4 Adaptation of Malaria Cell Atlas *P. falciparum* scRNA-seq

To adapt this dataset (from Dogga et al. 2024 [2]), the processed data (pf.zip) from the MCA website (https://www.malariacellatlas.org/downloads/pf.zip) was first downloaded. Transcripts from the same gene were summed, in order to match the bulk data used with this dataset. Cells identified as gametocytes were then removed, as these cells branch off into a different process of sexual development, and BLASE only supports single trajectories. Following this, 6000 cells were randomly selected from the remainder, to reduce the size of the final object.

Normalised counts were then calculated in the same manner as the *P. berghei* dataset, and Principal Component 1-3 and UMAP dimension 2-3 were used for visualisation.

Slingshot was used to calculate pseudotime (on the UMAP dimension, the high-resolution stage as the cluster labels, and “early ring” start cluster). TradeSeq’s `fitGAM` was then used with 7 knots, and the results of the association test were calculated in the same manner as the *P. berghei* dataset.

#### 6.6.5 Adaptation of spatial reference

To generate the scRNA-seq reference of Keratinocyte differentiation, pre-existing raw data in.bam format was downloaded from Wang et al. 2020 [21] (https://www.ncbi.nlm.nih.gov/geo/download/?acc=GSE147482&format=file).

Five samples are present. The following cutoffs were used for quality control:

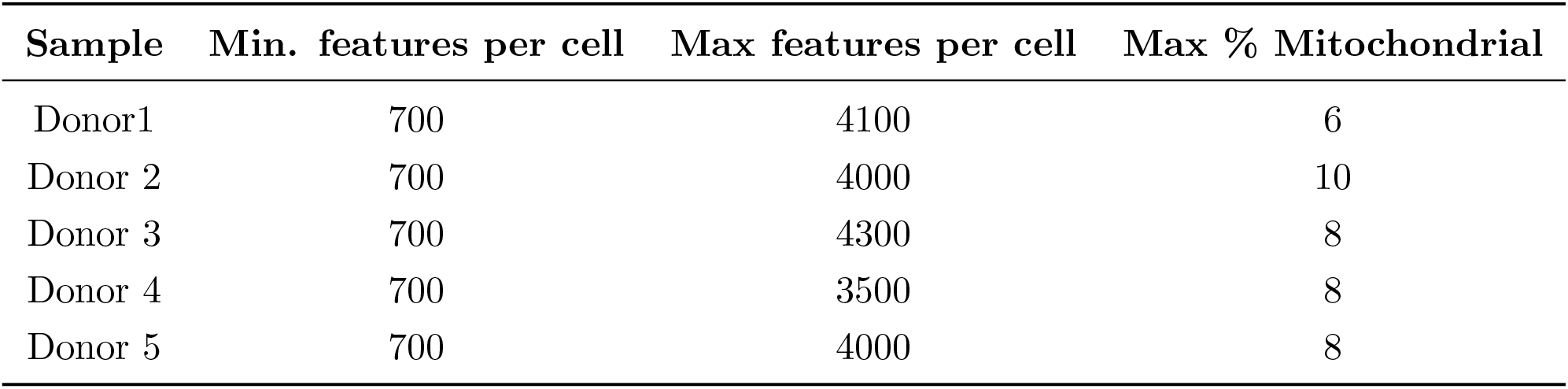

| Sample | Min. features per cell | Max features per cell | Max % Mitochondrial |
| --- | --- | --- | --- |
| Donor1 | 700 | 4100 | 6 |
| Donor 2 | 700 | 4000 | 10 |
| Donor 3 | 700 | 4300 | 8 |
| Donor 4 | 700 | 3500 | 8 |
| Donor 5 | 700 | 4000 | 8 |

Following quality control, Seurat was used to preprocess the data, using: `NormalizeData`, `FindVari-ableFeatures` (vst method, 2000 features), `ScaleData`, `RunPCA`. 14 PCs were selected by elbow plot. The samples were then integrated using STACAS unsupervised integration (1000 integration anchors). Seurat was then used to generate the KNN graph: `FindNeighbors` (on the PCA embedding), and finding clusters: `FindClusters` (resolution of 0.1). A UMAP embedding was then generated using `RunUMAP`.

A cluster of melanocytes were removed from the object, defined by high expression of the marker genes *MITF* and *MLANA* [21]. The remaining cells were then reprocessed, following the same steps as above, with 10 PCs, and a clustering resolution of 0.2. Basal (*KRT5, KRT14*), spinous (*KRT1, KRT10*), and granular (*DSC1, KRT2, SPINK5*) keratinocyte clusters were identified by high expression of their respective marker genes [21]. 6000 cells were randomly subset from this dataset, to reduce running time. Destiny [11] was then used to calculate pseudotime. The start index for destiny was found by taking the cell with the highest value in the first diffusion map dimension, determined to be the start of the trajectory by plotting the cell types in the diffusion map embeddings. To find genes variable over pseudotime, TradeSeq was used with 7 knots, only inspecting the highly variable genes identified earlier.

#### 6.6.6 Adaptation of MeDuSA scRNA-seq

This dataset, used to benchmark MeDuSA, was downloaded from the CytoTrace website (Examples page, “*C. elegans* ciliated neurons (10x)”, counts and metadata). Log normalized counts were generated using scran (`computeSumFactors`, default parameters) and bluster (`logNormCounts`, default parameters). Normalised counts were calculated as the exponential of the log normalised counts. The “Ground truth” metadata was used as pseudotime. TradeSeq was then run with 8 knots (selected in the same way as the other datasets).

### 6.7 Generation of bulk RNAseq datasets

A copy of each of these objects, and code to reproduce them is provided on Zenodo (subsection 7.1).

#### 6.7.1 Adaptation of Zhang et al. 2021 *P. falciparum* heat shock Piggybac knockout RNA-seq

To prepare this dataset for use with BLASE, data from the original paper’s (Zhang et al. 2021 [33]) Supplementary Data 3 spreadsheet was downloaded (https://static-content.springer.com/esm/art%3A10.1038%2Fs41467-021-24814-1/MediaObjects/41467_2021_24814_MOESM5_ESM.xlsx). FPKM counts were extracted and used these as normalised counts. Transcripts from the same gene were collapsed by summing of their counts to maintain consistency with the scRNA-seq that would be used with this data.

#### 6.7.2 Adaptation of Otto et al. 2014 *P. berghei* RNA-seq

To prepare this dataset for use with BLASE, the spreadsheet with FPKM reads from the original paper’s (Otto et al. 2014 [54]) Additional file 9 was downloaded (https://static-content.springer.com/esm/art%3A10.1186%2Fs12915-014-0086-0/MediaObjects/12915_2014_86_MOESM9_ESM.xlsx), selecting only the *P. berghei* mRNA abundance data.

#### 6.7.3 Adaptation of Ganier healthy skin Visium

The original data (Ganier et al. 2024 [3]) for several healthy skin Visium runs was downloaded from the Spatial Skin Atlas (https://spatial-skin-atlas.cellgeni.sanger.ac.uk/). Greyscale histology images were generated in Python. These data files were loaded by Seurat. Majority keratinocyte spots were then selected for use with BLASE.

The proportion of keratinocytes in each spot was calculated from the results of cell2location included in the objects as downloaded. Proportions were considered keratinocytes if they were in either the “Basal keratinocytes” or “Suprabasal keratinocytes” clusters.

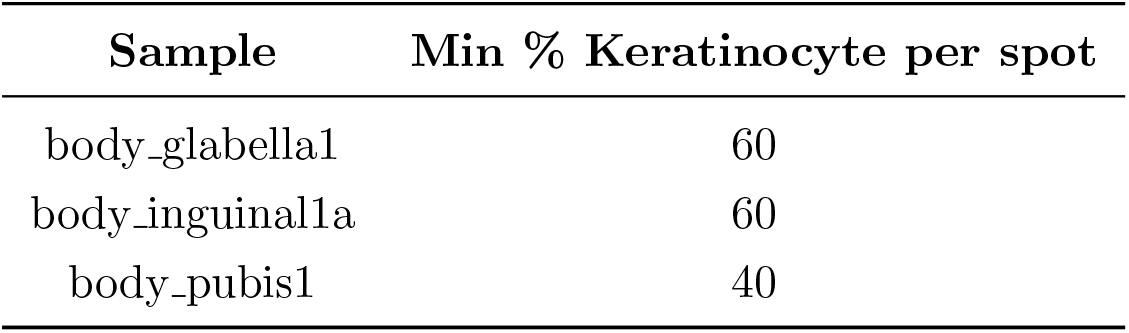

| Sample | Min % Keratinocyte per spot |
| --- | --- |
| body_glabella1 | 60 |
| body_inguinal1a | 60 |
| body_pubis1 | 40 |

Normalised counts for these data were calculated by taking the exponential of the log counts provided in the data.

#### 6.7.4 Adaptation of Ma psoriatic skin Visium

Visium data for skin with psoriasis was downloaded from the Gene Expression Omnibus (https://www.ncbi.nlm.nih.gov/geo/query/acc.cgi?acc=GSE225475), originally generated by Ma et al. in 2023 [20]. Once again, greyscale histology images were generated in Python, and these data files were loaded by Seurat.

To identify spots likely to be high in keratinocytes, spots with high expressions of *KRT5* and *KRT10* were selected. Normalised counts were again given by the exponential of the log counts in the objects.

| Sample | Min. log KRT5 | Min. log KRT10 |
| --- | --- | --- |
| PP1 | 3 | 3 |
| PP2 | 3 | 3 |
| PP3 | 3 | 3 |
| PP4 | 3 | 3 |

#### 6.7.5 Adaptation of Painter et al. 2018 *P. falciparum* Microarray

To adapt this dataset [26] for this analysis, the data was downloaded from https://static-content.springer.com/esm/art%3A10.1038%2Fs41467-018-04966-3/MediaObjects/41467_2018_4966_MOESM4_ESM.xlsx, and the “Estimated Total Abundance” sheet was used. Gene names were changed to use “-” instead of “ “.

#### 6.7.6 Adaptation of López-Barragán 2011 *P. falciparum* RNA-seq

To use this dataset [29], it was downloaded from PlasmoDB [55] using this search: https://plasmodb.org/plasmo/app/workspace/strategies/import/1a7322955d116b6b. “.1” was removed from transcript names. Only unique reads were saved for analysis.

#### 6.7.7 Adaptation of Otto et al. 2010 *P. falciparum* RNA-seq

This dataset was downloaded from PlasmoDB using this search: https://plasmodb.org/plasmo/app/workspace/strategies/import/698e68176373afcb. “.1” was removed from transcript names. Only unique reads were saved for analysis.

### 6.8 Pseudobulking of scRNA-seq datasets

Datasets presented here were pseuodbulked by pseudotime bin assigned by BLASE.

Cells were randomly split into two equally sized groups, one of which remains as a scRNA-seq reference, and one as pseudobulk samples. When pseudobulking by cell-type annotation, the normalised reads for each annotation were summed, giving one value for each gene per annotation.

For the simulated testing dataset, cells were randomly assigned to one of two replicates for each pseudotime bin. The `get bins as bulk` function from BLASE summed the normalised counts of genes for each bin-replicate pair to give two replicate pseudobulk samples per bin. A cutoff of at least 20 cells per bulk was used. For bins with enough cells to meet this minimum in a given replicate, the cells are split by their assigned replicate. Should there not be enough cells in a bin, the `get bins as bulk` function will randomly sample 75% of a bin twice to generate the two replicates, if this will provide enough cells to meet the minimum cells per bulk cutoff, otherwise a warning is given and only one replicate is produced. To pseudobulk the haematopoiesis dataset, the cells were split into evenly sized test and train groups, and in the test group, the expression of all cells in each pseudotime bin was summed to generate the pseudobulk data.

### 6.9 Application of BLASE to datasets

Due to each reference dataset having individual differences due to their species, cell types and trajectories, different parameters were required for optimal performance. BLASE’s hyperparameter tuning and geneselection methods provide heuristics to select the number of bins and genes to be used in the mapping. A comprehensive list of the parameters used when benchmarking BLASE can be seen in Table A.3 and Table A.4. Exact code to reproduce results is available on Zenodo (Availability of data and materials).

### 6.10 Downstream analysis

#### 6.10.1 Comparable differential expression analysis of bulk and scRNA-seq datasets

Differential expression (DE) analysis was performed using the limma R package [56]. The FPKM adjusted counts were passed to limma, and were processed using: `lmFit`, `contrasts.fit` and `eBayes` with the default parameters. The contrast matrix was generated by creating a group for each combination of pairs from [strain] (“NF54”, “PB31”, “PB4”) and [growth condition] (“Normal” or “HS”).

For the single cell data, raw pseudobulked counts for the bins being compared were first normalised using limma’s `voom` function (default parameters).

#### 6.10.2 Differential expression analysis of scRNA-seq datasets

To find DE genes from scRNA-seq data (for example to find genes for GO terms in Additional file 2), the `findMarkers` function in Seurat was used with default parameters between two groups. The results were then filtered by “p_val_adj” ¡ 0.05 and “avg_log2FC” ¿ 0.5.

#### 6.10.3 Gene ontology term enrichment analysis

GO term enrichment was carried out using the topGO R package [57]. Gene GO term mappings were taken from PlasmoDB v68 for Pf3D7 [55]. To generate the exact format required by TopGo, code from the function `formatGOdb` in PfGO [58] was adapted. topGO objects were created for each comparison, using the “BP” ontology, *allGenes* the entire set of genes in *P. falciparum*, and the selected genes were every significantly (adjusted P value *<* 0.05) differentially expressed gene. `runTest` was executed with the “classic” algorithm and “fisher” statistic.

### 6.11 Application of existing deconvolution methods to datasets

Several existing deconvolution methods were tested alongside BLASE: CIBERSORTx, DWLS, MeDuSA, and MuSiC [5, 7, 8, 42, 9].

The sections below contain details of the default parameters (and exceptions to these) used for comparing each tool with BLASE.

#### 6.11.1 CIBERSORTx

CIBERSORTx was run as a docker container. Before use, all gene, cell and sample names had hyphens (-) and underscores () removed. Normalised counts were used. The maximum genes to use (G.max) was set to *min*(500, *length*(*genes*(*singleCellExperiment*) − 100. The number of permutations (perm) was set to 200.

#### 6.11.2 DWLS

All gene names had hyphens (-) replaced with underscores (). Raw and log counts were used to create a Seurat v5 object [48]. A signature matrix was constructed with `buildSignatureMatrixUsingSeurat` (diff.cutoff=0.5, pval.cutoff=0.05).

Then, each bulk sample was iteratively deconvoluted, using `trimData` (default parameters), and `solveDampenedWLS` (default parameters). Pseudotime bins with a returned proportion of *<* 0 were set to 0.

#### 6.11.3 MeDuSA

All gene names has underscores () replaced with hyphens (-). A Seurat v5 object [48] was then created using raw and log counts. The `MeDuSA` function was then run with:

- markerGene = NULL
- span = see Table 2
- resolution = see Table 2
- smooth = TRUE
- fractional = TRUE
- nbins = see Table 2

**Table 2:** Details of the parameters used for application of MeDuSA to datasets.

| Dataset | resolution | span | nbins |
| --- | --- | --- | --- |
| Simulated | 5 | 2 | 4 |
| Myeloid Differentiation | 5 | 2 | 5 |
| *P. falciparum* lifecycle | 6 | 2 | 6 |
| *P. falciparum* heat shock | 6 | 2 | 6 |
| *C. elegans* MeDuSA reproduction | 5 | 2 | 5 |
| *C. elegans* MeDuSA reproduction | 10 | 2 | 5 |
| *C. elegans* MeDuSA reproduction | 5 | 4 | 5 |
| *C. elegans* MeDuSA reproduction | 5 | 1 | 5 |
| *C. elegans* MeDuSA reproduction | 5 | 2 | 10 |
| *C. elegans* MeDuSA reproduction | 5 | 2 | 2 |

Specific datasets used different parameters for span, resolution and nbins for MeDuSA, which are shown in Table 2.

Following this, `MeDuSA_VarExplain` was run with default parameters. Note that a smaller value for span may reduce the wide spread in the results, however, when set to a low value the function fails to complete, warning that the span is set too low.

#### 6.11.4 MuSiC

MuSiC was used with default parameters, using the `music prop` function.

### 6.12 Benchmarking runtime of tools

Benchmarking of BLASE’s runtime was done following the R pseudocode in Algorithm 4.

The benchmarking was run on a server with:

#### Algorithm 4

R code used for timing deconvolution time

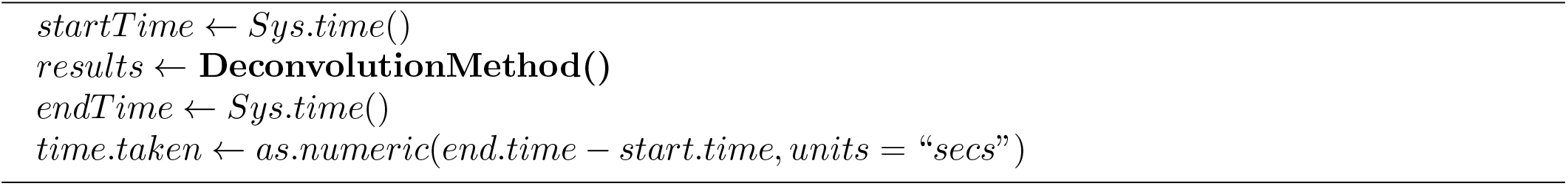

- CPU - 4x Intel(R) Xeon(R) CPU E7-4890 v2 @ 2.80GHz
- Memory - PC-12800/1600Mhz - 96 x 32Gb modules
- OS - Ubuntu 22.04.3 LTS

When benchmarking CIBERSORTx, the input data was written to disk before timing began, to create as fair a comparison as possible.

## 7 Declarations

### 7.1 Availability of data and materials

BLASE Source Code available under the GPLv3 license on GitHub at https://github.com/andrewmccluskey-uog/blase

BLASE Documentation available at https://andrewmccluskey-uog.github.io/blase/

The datasets generated and/or analysed during the current study are available in the “BLASE Reproducibility Scripts & Data” repository on Zenodo, at https://doi.org/10.5281/zenodo.16615702

BLASE Package will be available for download from Bioconductor

### 7.2 Authors’ Contributions

**A.M**.: Methodology, Software, Formal analysis, Investigation, Data Curation, Writing - Original Draft, Visualization **T.K**.: Methodology, Writing - Original Draft **A.M.S**.: Conceptualization, Funding Acquisition, Supervision, Writing - Review & Editing **R.K**.: Supervision, Writing - Review and Editing **D.A.G**.: Conceptualization, Funding Acquisition, Writing - Review & Editing **T.D.O**.: Supervision, Conceptualization, Resources, Project administration, Funding acquisition, Writing - Original Draft

## 7.3 Acknowledgements

We would like to thank: Surya Koturan, Joy Kabagenyi, Elcid Aaron Pangilinan, Yiyi Cheng and Anna Bachmann for feedback on the manuscript and tool.

## 7.4 Ethics approval and consent to participate

Not Applicable

### 7.6 Datasets

All objects and scripts to generate the results described above are available in Zenodo under DOI: 10.5281/zenodo.16615703

### 7.7 Funding

**A.M**. was funded by the Biotechnology and Biological Sciences Research Council CASE studentship [BB/X511389/1] and Unilever. **T.K**. was funded by Engineering and Physical Sciences Research Council [EP/T517896/1]. **A.M.S**., **R.K. and D.A.G**. are employed by Unilever. **T.D.O**. was supported by the Wellcome Trust [104111/Z/14/Z&A] and the ExposUM Institute of the University of Montpellier [grants ANR-21-EXES-0005 and Occitanie Region].

## 8

**Bibliography**

## Appendix A

### Supplemental

**Figure A.1:**
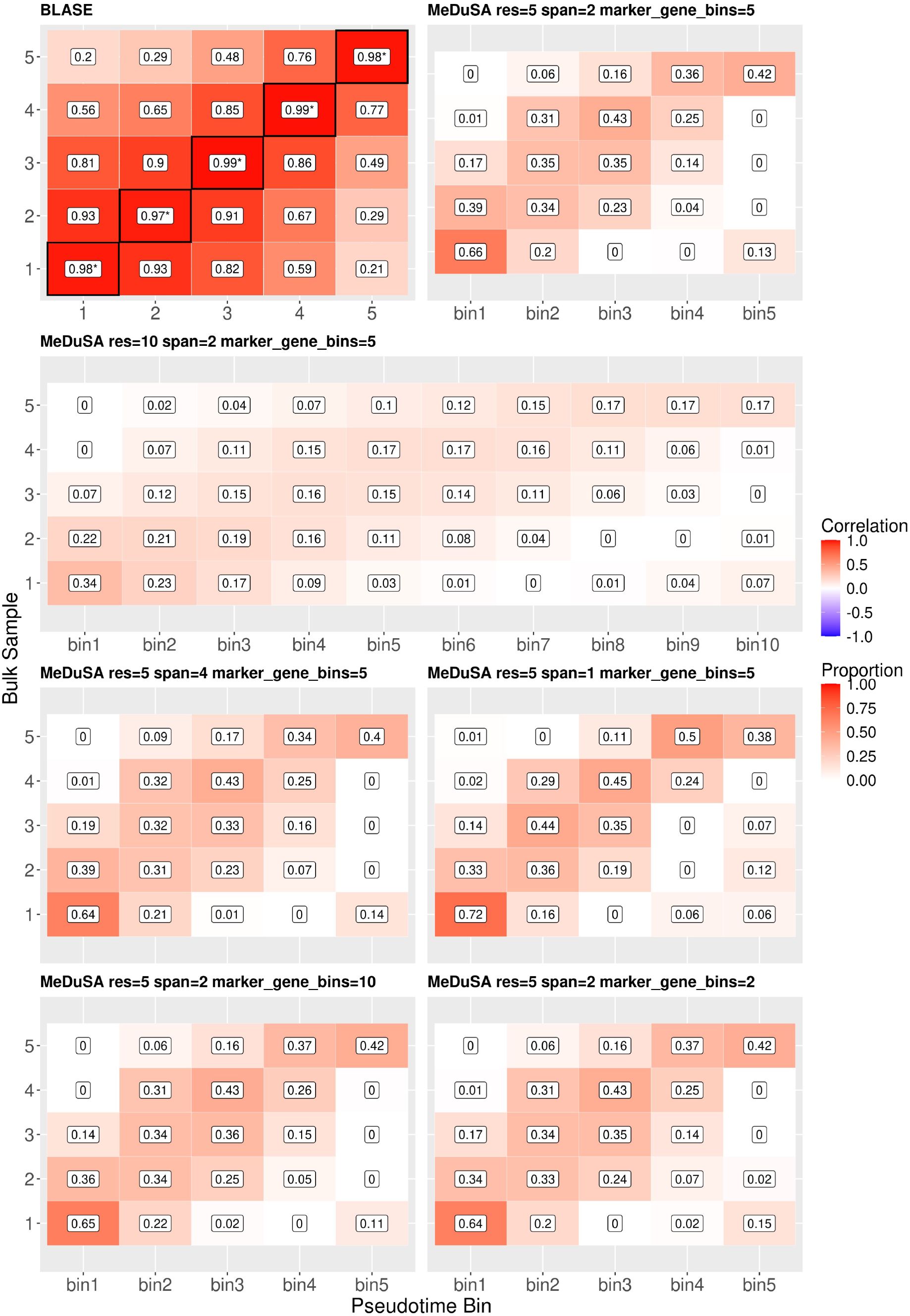
Plots showing results of BLASE and MeDuSA with varying parameter choices from the *C. elegans* ciliated neurons dataset used to benchmark MeDuSA by its authors.

**Figure A.2:**
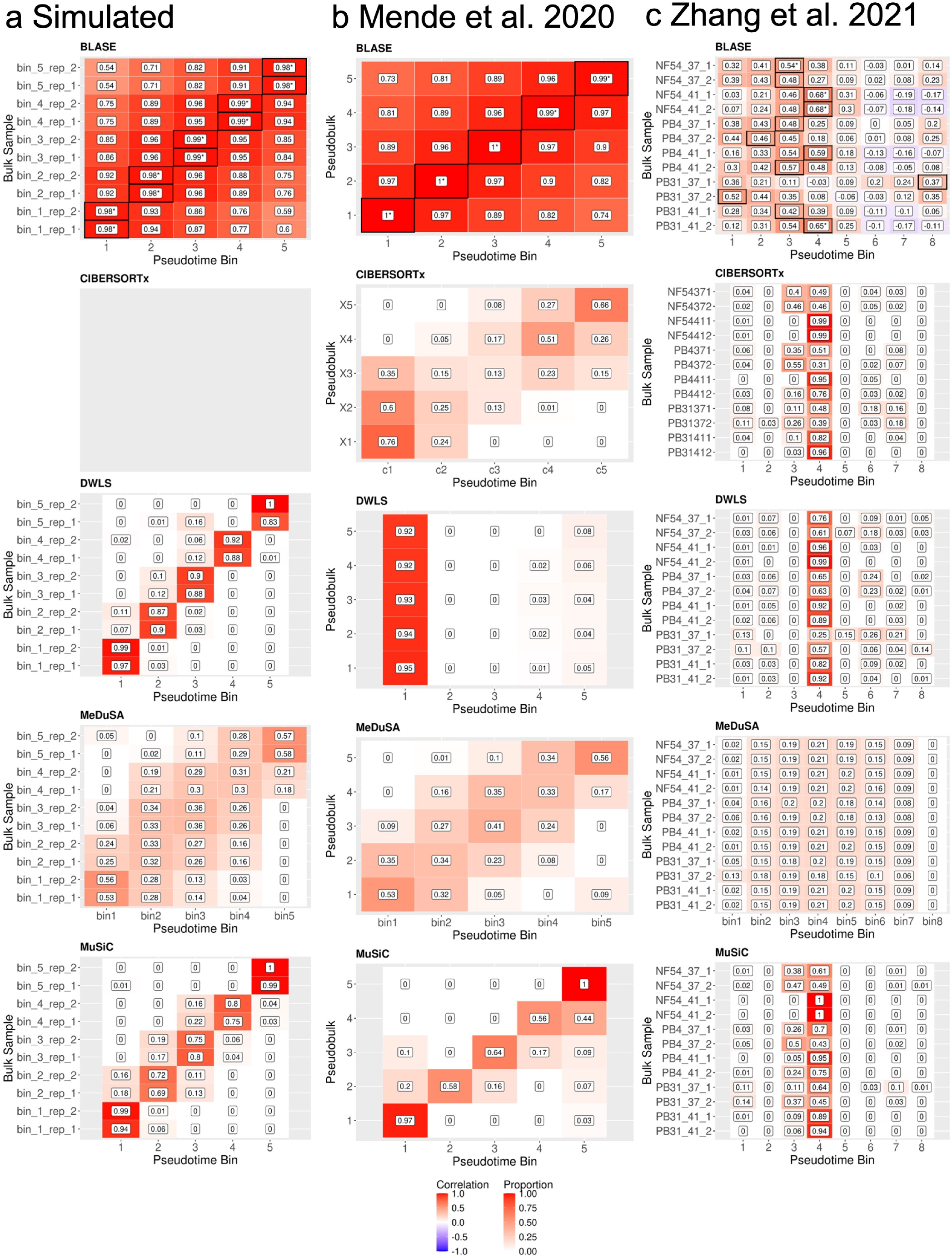
BLASE performance compared with other tools on several datasets a, BLASE (All genes), CIBERSORTx, MuSiC, MeDuSA, and SCDC result heatmaps showing the predicted mappings on a simulated dataset (using a test/train split). b, The results from running the same tools as a on the Mende et al. 2020 haematopoiesis dataset (test/train split). BLASE used a TradeSeq generated genelist. c, The results from running the same tools as a on the Dogga et al. 2023 scRNA-seq with the Zhang et al. 2021 bulk RNA-seq data. BLASE used a Gene Peakedness gene selection.

**Figure A.3:**
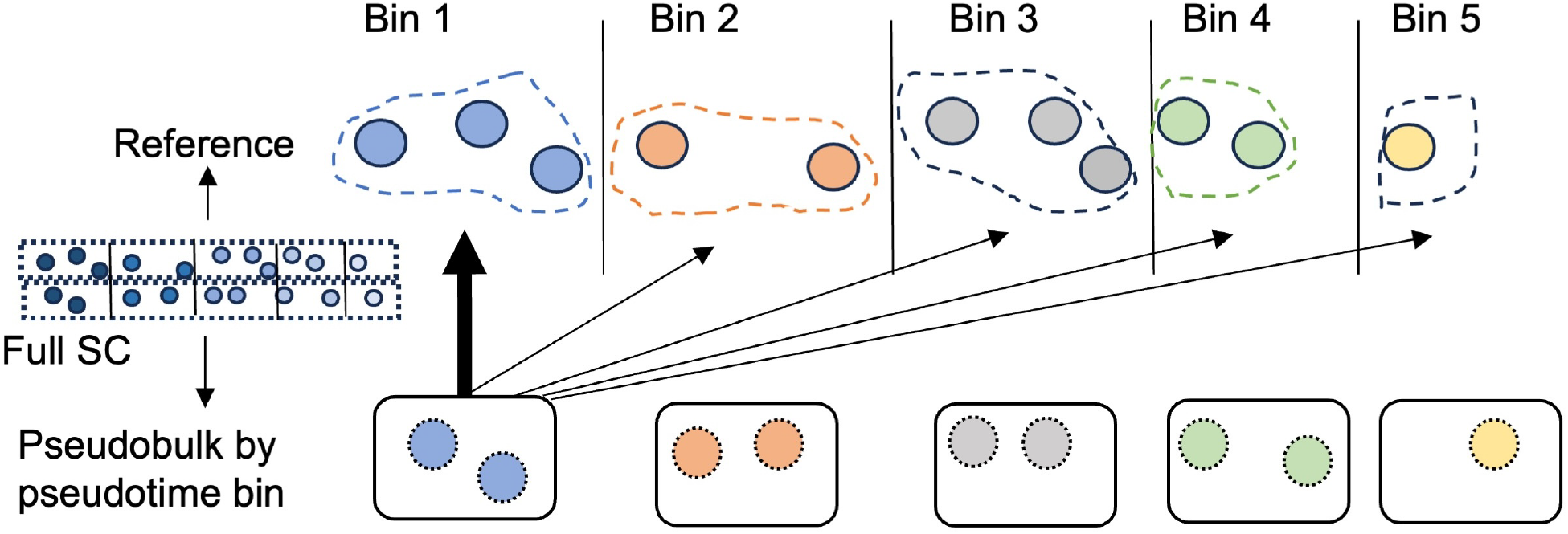
Generation of testing datasets. A scRNA-seq dataset was generated, then randomly split into two groups. One group was used as reference, and the second group was pseudobulked by pseudotime bin.

**Figure A.4:**
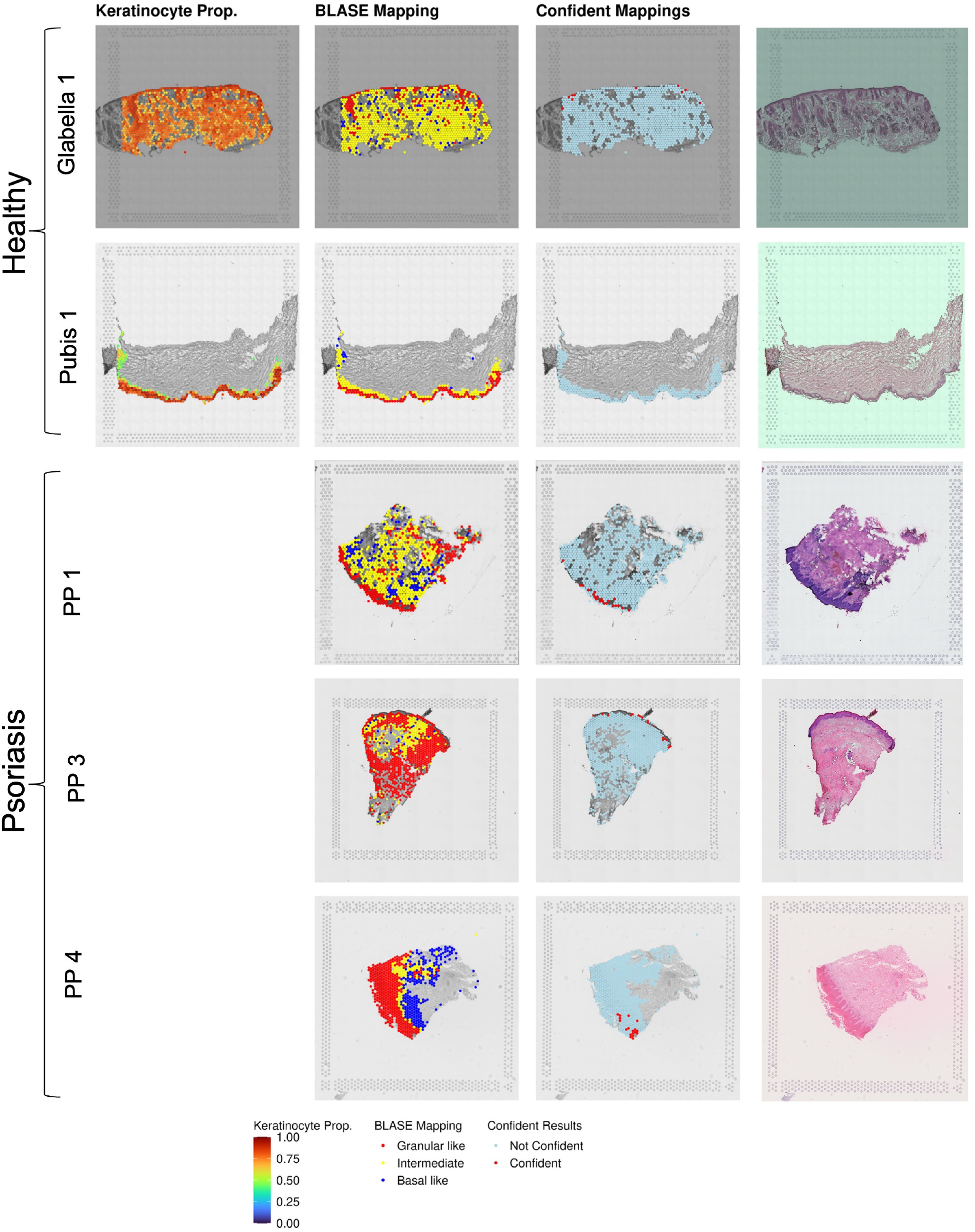
Plots showing the following for a wider selection of Visium skin samples from Ganier et al. 2024 (healthy) and Ma et al. 2023 (psoriasis): Keratinocyte proportion in spot (Ganier et al. 2024 only), BLASE mappings for keratinocyte maturity in each spot, the spots with a confident BLASE mapping, and the original H&E stained samples on the Visium Slide.

**Figure A.5:**
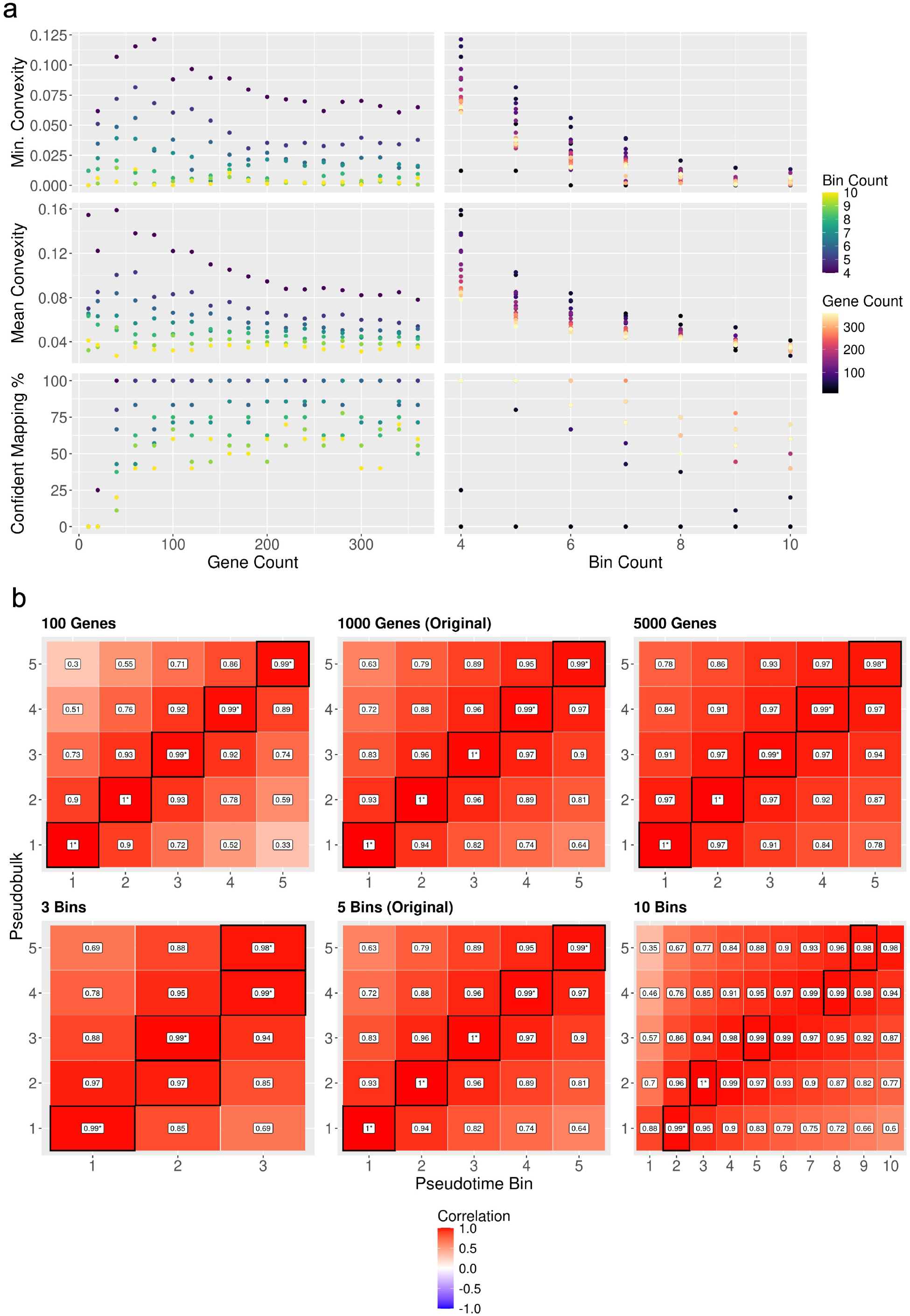
a, BLASE Hyperparameter selection plot, created with `find_best_params_results()`. This plot shows the worst and mean “convexity” of a set of parameters. Convexity is calculated by pseudobulking each bin, and using BLASE to map them, and finding the distance between the best mapping (in the expected case, the bin the pseudobulked sample is generated from) and the bin with the second highest correlation. A higher “convexity” is better. The plots show the x-axis as either the number of genes used (left side) or the number of bins used (right side). Also shown is the percentage of confident mappings for each pair of n bins and genes. b, A set of BLASE result heatmaps showing the effect of varying numbers of genes and bins on the same dataset. Results generated from the Mende et al. 2020 haematopoiesis scRNA-seq dataset.

**Figure A.6:**
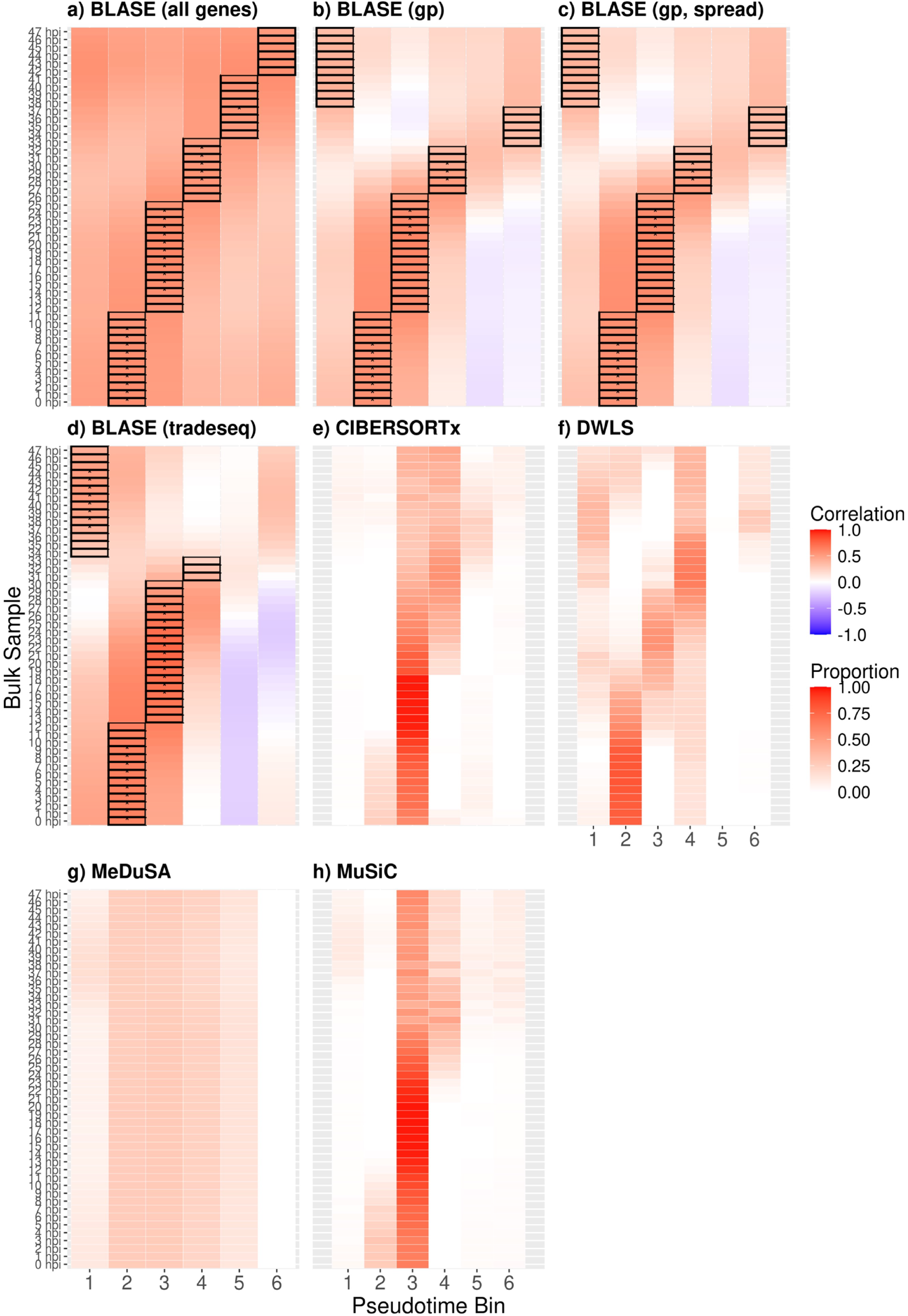
BLASE gene selection methods compared with deconvolution packages (Painter et al. 2018 / Dogga et al. 2023) a, BLASE result heatmap showing the predicted mappings in the Dogga et al. 2023 scRNA-seq for bulk RNA-seq the from Painter et al. 2018, using all genes. b, BLASE result heatmap showing the same data as in a, when genes selected by the BLASE gene peakedness method. c, BLASE result heatmap showing the same data as in a, when genes selected by the BLASE gene peakedness spread method. d, BLASE result heatmap showing the same data as in a, when genes selected by the TradeSeq associationTest. e, Cell proportion estimates predicted by CIBERSORTx for the datasets described in a. f, Cell proportion estimates predicted by DWLS for the datasets described in a. g, Cell proportion estimates predicted by MeDuSA for the datasets described in a. h, Cell proportion estimates predicted by MuSiC for the datasets described in a.

**Figure A.7:**
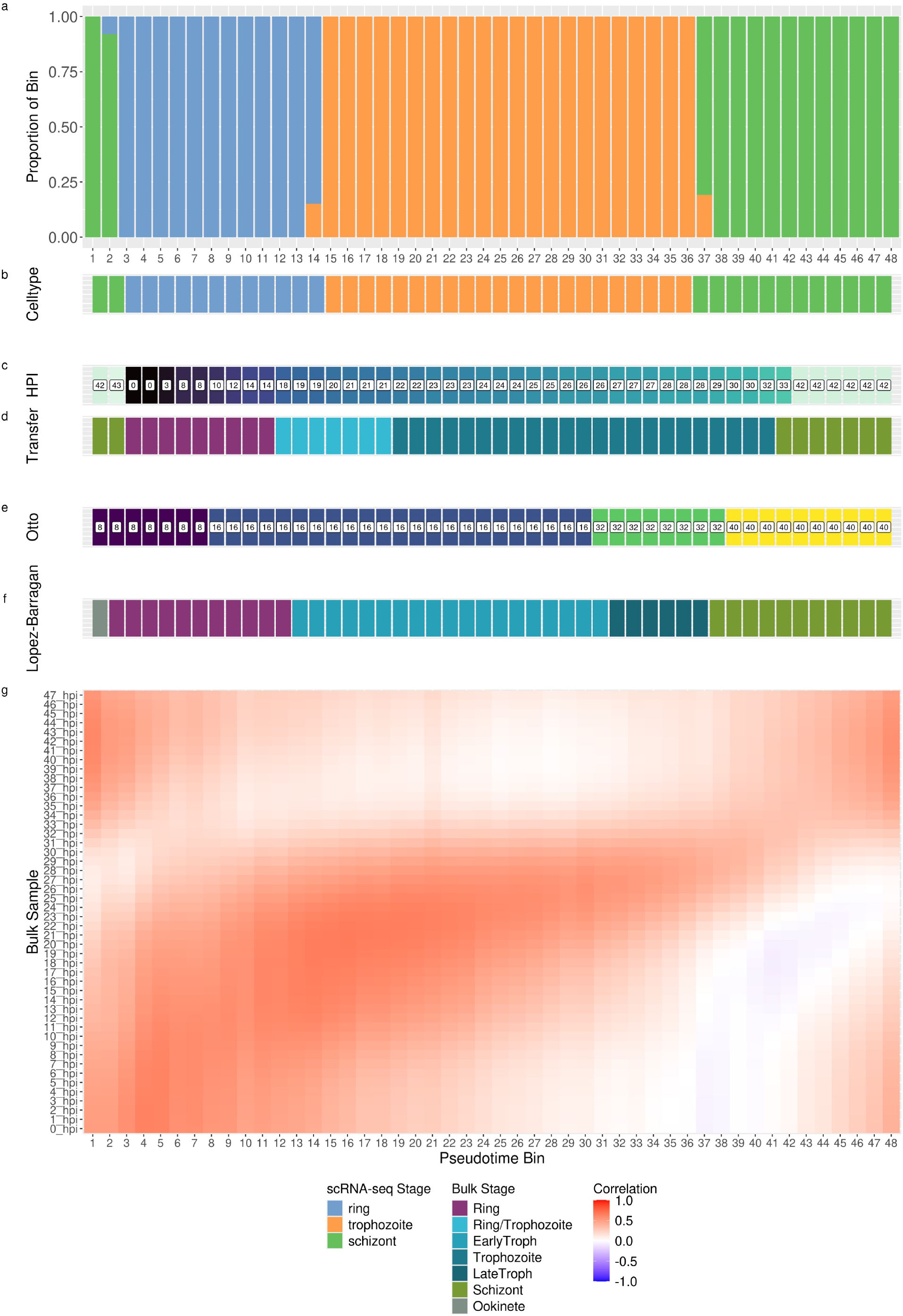
Annotating *P. falciparum* scRNA-seq from bulk RNA-seq. a, Pseudotime bin populations annotated by original scRNA-seq analysis. b, Major celltype for each pseudotime bin as annotated in the original scRNA-seq (i.e. cell type with the highest proportion) c, The bulk sample (hours post infection, HPI) from Painter et al. that BLASE predicts the pseudotime bin is most likely to be from. d, The bulk sample from c, but with the expected cell types from the bulk data annotated. e, The same as c, but mapping bins to Otto et al. data. f, The same as c, but mapping bins to López-Barragán et al. data. g, The BLASE Mapping heatmap of the data used to calculate this, with 48 bins split by cell.

**Figure A.8:**
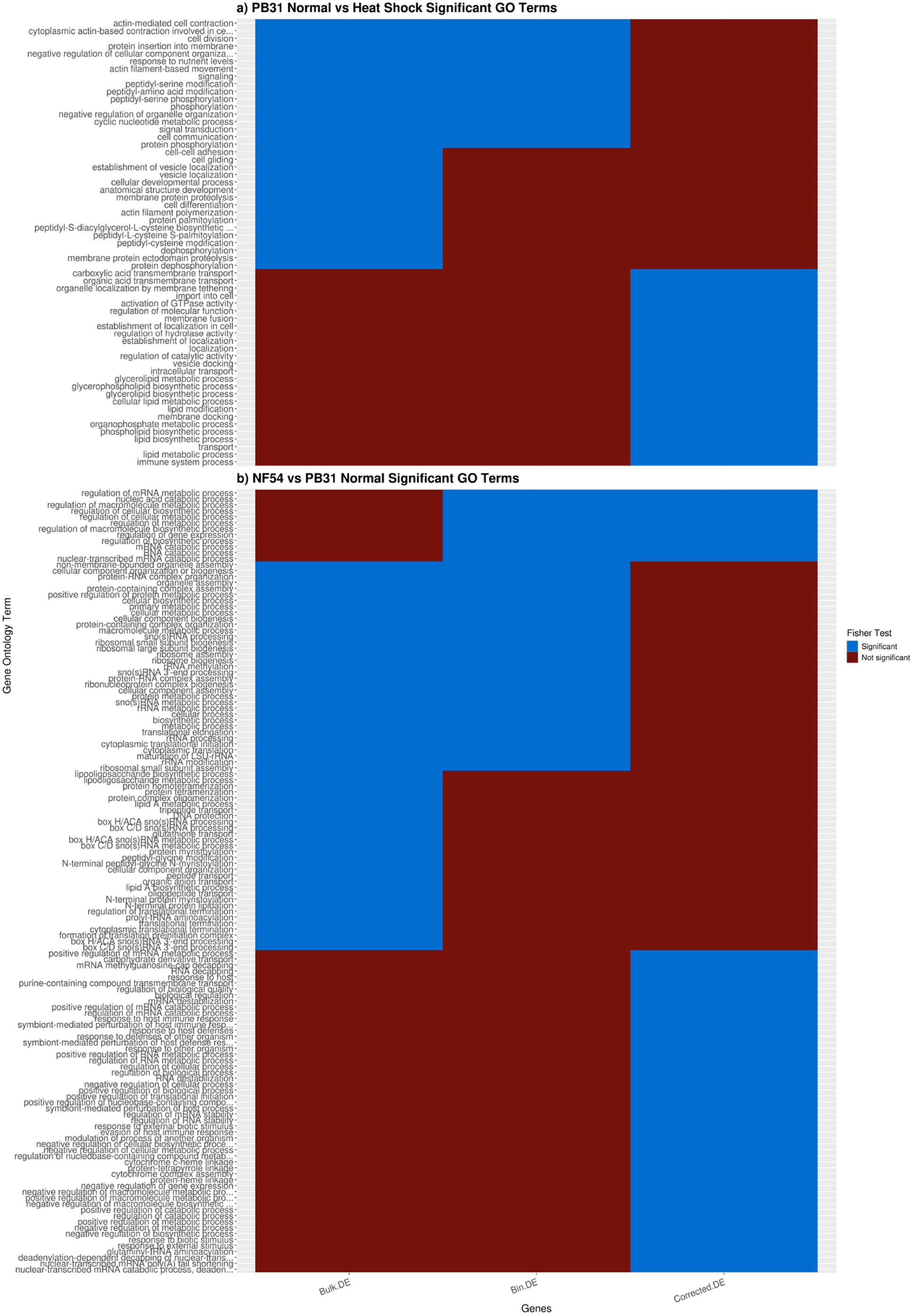
Comparison of Fisher Statistic for GO enrichment analysis on gene lists from uncorrected bulk DE genes, BLASE corrected DE genes, and the list of removed genes for three comparisons: a,, NF54 vs. PB31 in normal conditions, and b, PB31 in normal vs. heat shock conditions (note, no corrections were made for the NF54 normal growth conditions vs. heat shock conditions comparison, so there was no difference in GO term enrichment analysis).

**Figure A.9:**
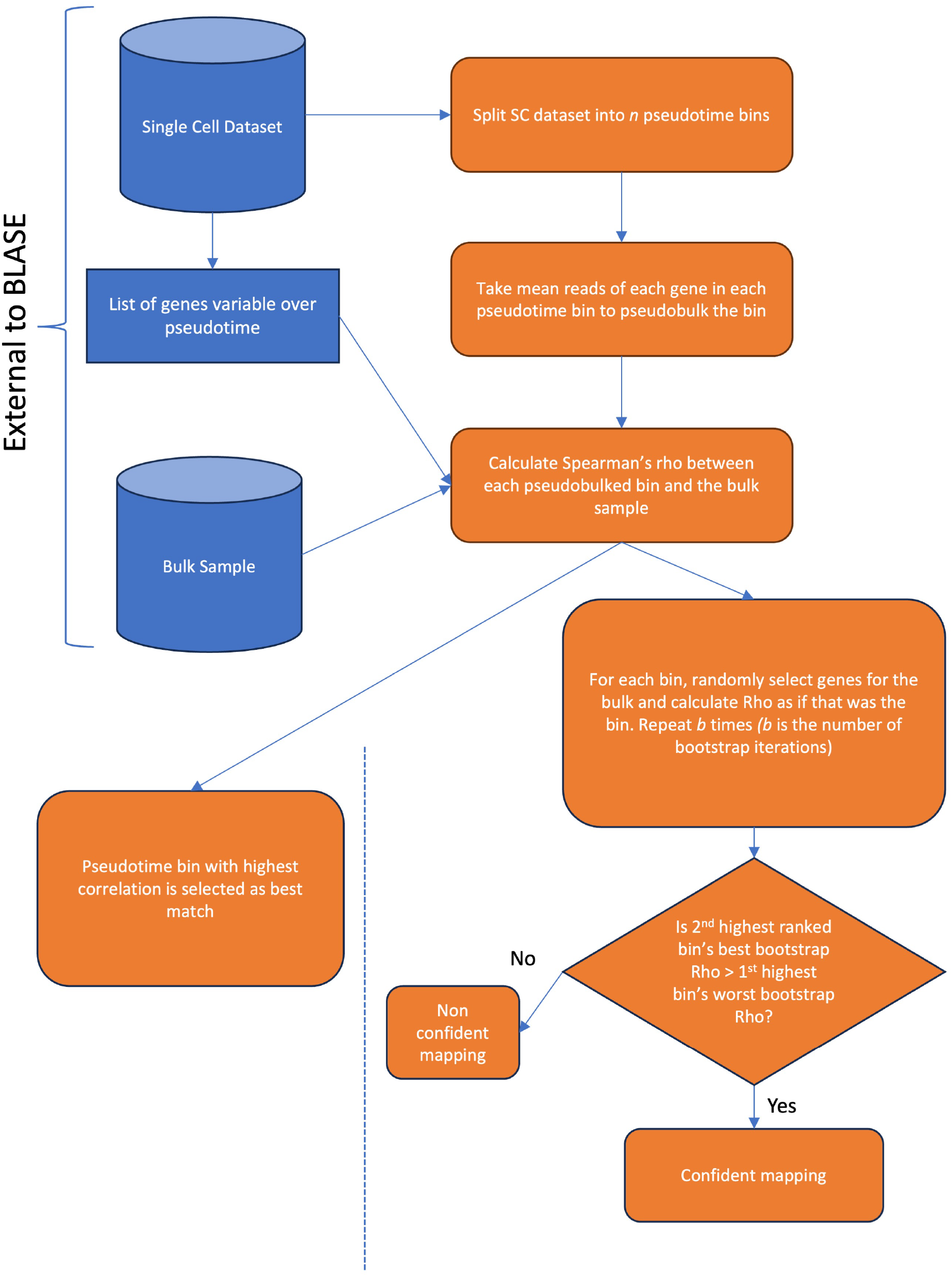
Flowchart detailing the steps of the BLASE algorithm. Three inputs (scRNA-seq reference, list of genes that change over pseudotime, and a bulk RNA-seq sample to map) are shown in blue. Steps taken by BLASE are shown in orange.

**Figure A.10:**
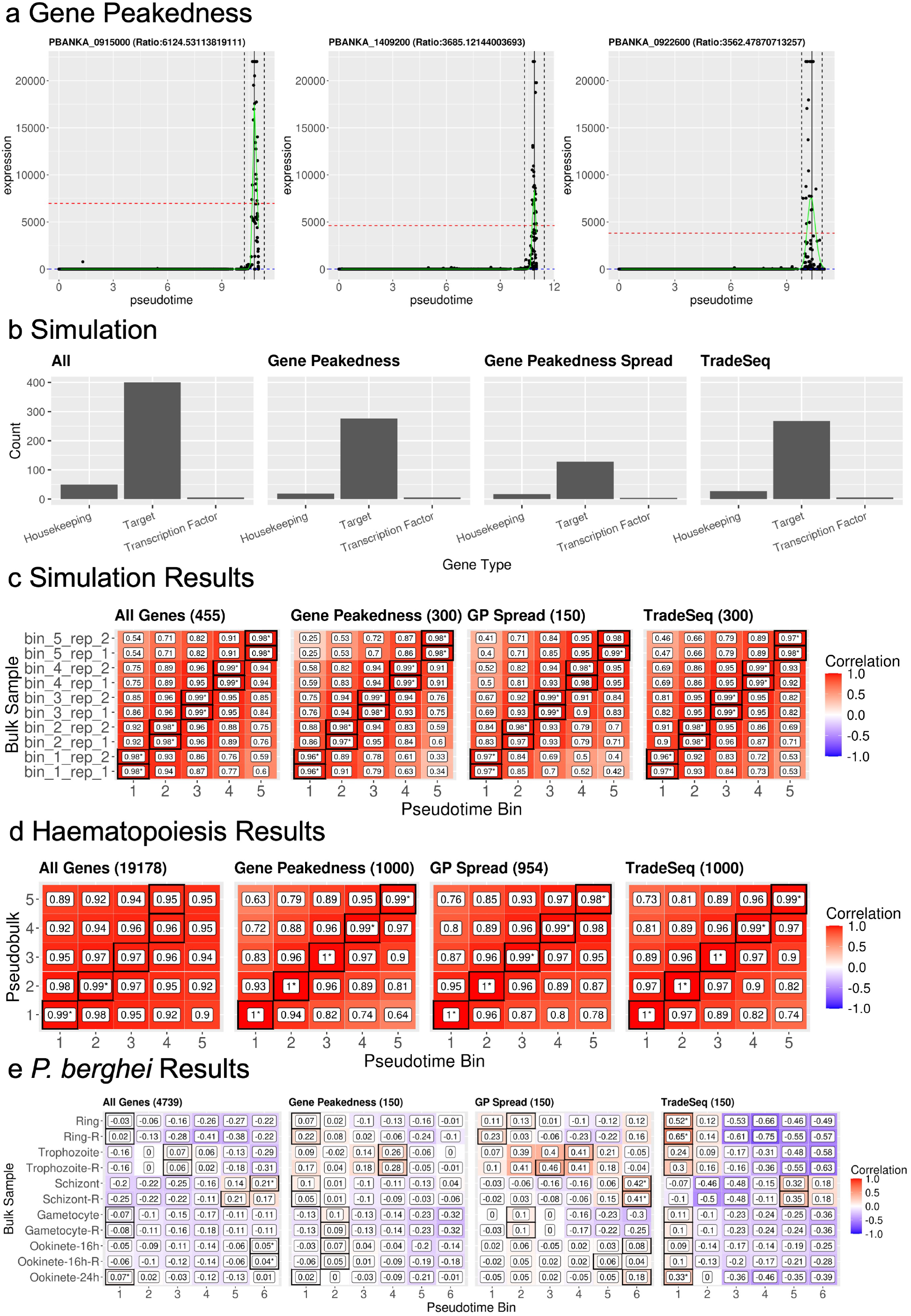
Gene selection methods for BLASE a, The top three genes (sorted by descending BLASE gene peakedness) for the Howick et al. 2019 *P. berghei* reference dataset. Normalised counts are plotted as a scatter plot. Two dotted black lines show the 10% window around the peak, as calculated by the smoothed values from a GLM fitted to the gene. The dashed red line indicates the mean count within the window, and the dashed blue line the mean outside the window. b, Counts of the types of simulated gene in the simulated test dataset, for all genes, the genes selected by tradeSeq, the genes selected by BLASE gene peakedness and the genes selected by BLASE spread gene selection. These methods filter out housekeeping genes which are not informative about the trajectory. c,d,e, BLASE result heatmaps generated using genes selected by four methods (all genes, BLASE gene peakedness selection, gene peakedness spread, tradeSeq selection), for three scRNAseq references. c: Simulated dataset, d: Mende et al. 2020 haematopoiesis dataset, e: Howick et al. 2019 *P. berghei* dataset.

**Figure A.11:**
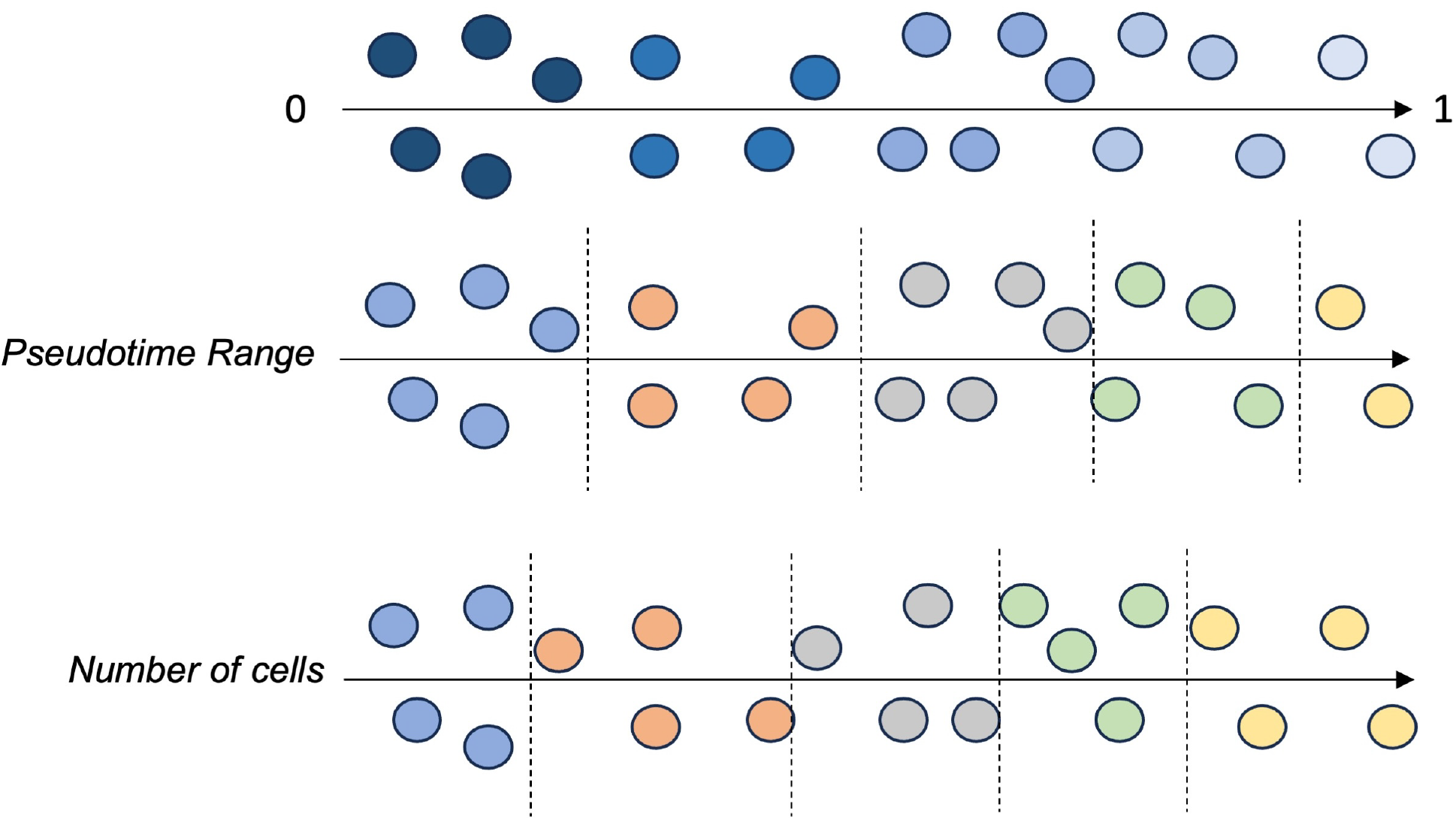
Schematics of BLASE pseudotime bin selection methods. Pseudotime range: ensures all bins have the same width of pseudotime (in this case, 0.2). This allows varying numbers of cells. This is the recommended method as it preserves a constant transcriptional “distance.” Cells: ensures all bins have the same number of cells, allowing varying widths of pseudotime.

**Table A.1:**
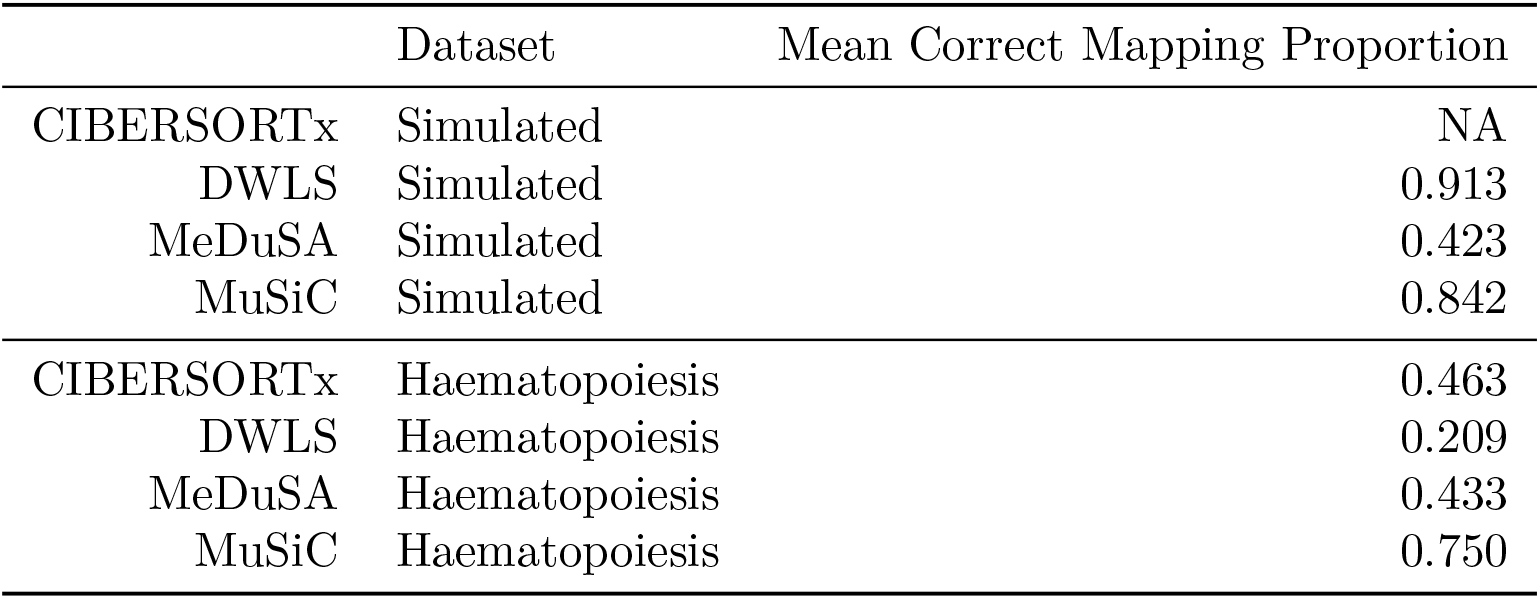
Mean correctly mapped proportions for other deconvolution tools, rounded to 3 decimal places.

|  | Dataset | Mean Correct Mapping Proportion |
| --- | --- | --- |
| CIBERSORTx | Simulated | NA |
| DWLS | Simulated | 0.913 |
| MeDuSA | Simulated | 0.423 |
| MuSiC | Simulated | 0.842 |
| CIBERSORTx | Haematopoiesis | 0.463 |
| DWLS | Haematopoiesis | 0.209 |
| MeDuSA | Haematopoiesis | 0.433 |
| MuSiC | Haematopoiesis | 0.750 |

**Table A.2:** *P. falciparum* intraerythrocytic lifecycle stages, and their mappings in two datasets (Painter et al. 2018, Dogga et al. 2023). Ranges are inclusive. 12 pseudotime bins were used.

| Stage | HPI (Painter et al.) | Pseudotime Bins with $> 1$ cell (Dogga et al.) |
| --- | --- | --- |
| Ring | 0-21 | 1-3 |
| Trophozoite | 16-32 | 3-4 |
| Schizont | 33-48 | 2-6,1 |

**Table A.3:**
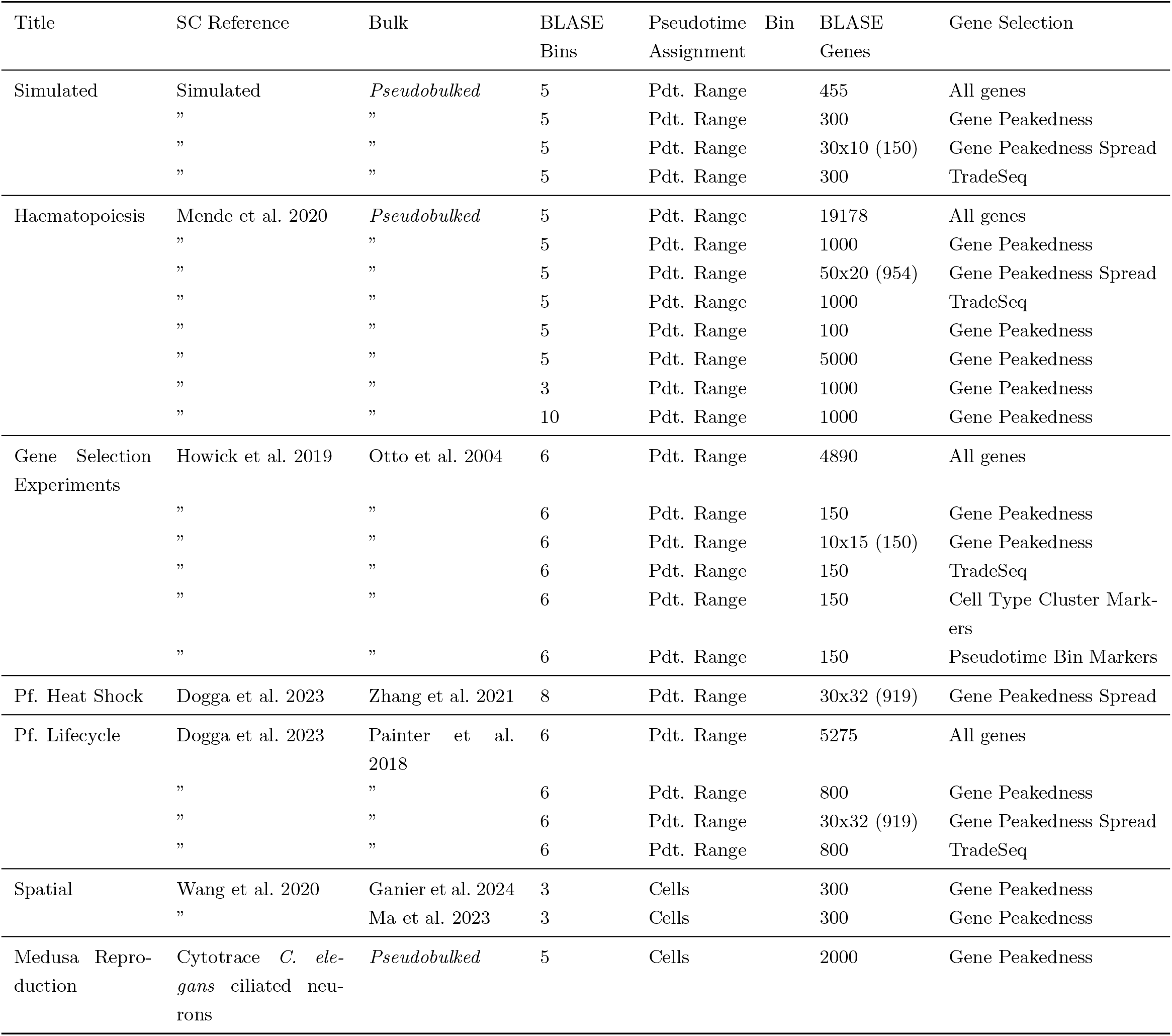
A comprehensive list of the BLASE hyperparameters used to generate the results presented in this paper. Pseudotime is abbreviated as “Pdt.”.

**Table A.4:**
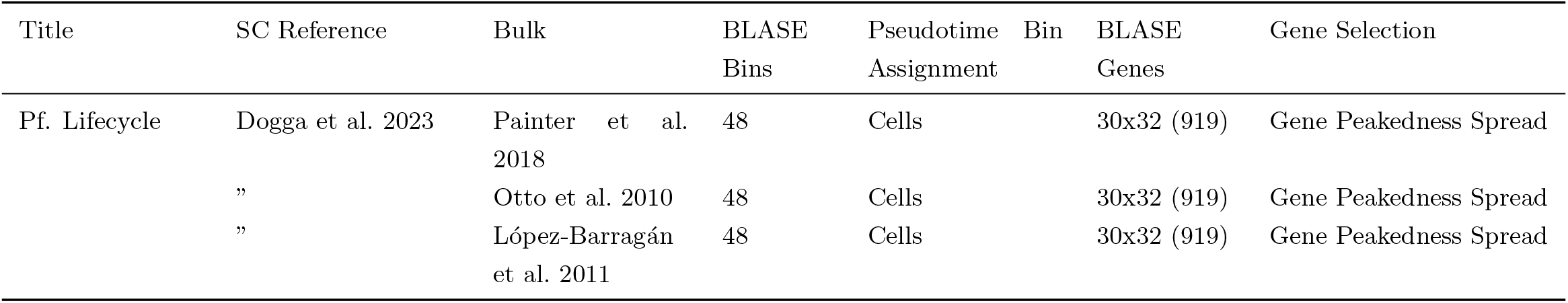
A list of the BLASE hyperparameters used to annotate scRNA-seq psuedotime bins with known bulk RNA-seq as presented in this paper. Pseudotime is abbreviated as “Pdt.”.

## Appendix B

### Additional Files

#### B.1 Additional file 1

**Filename**: SF1_Zhang_PF_bin1_vs_bin4_sc_GO_terms

**Format**: CSV

**Description**: All significantly enriched GO terms between Bin 1 and Bin 4, based on scRNA-seq DEGs.

#### B.2 Additional file 2

**Filename**: SF2_Zhang_PF_PB31_normal_v_HS_bulk_and_sc_de_genes

**Format**: CSV

**Description**: Table of the DE genes with ordered by lowest adjusted p-value, between the bulk samples, and between the pseudobulked pseudotime bins which were the best matches for the bulk RNA-seq samples in the comparison of PB31 normal conditions against heat shock conditions. “logFC” columns are rounded to 1 decimal place. “adj.P.Val” columns are rounded to 3 decimal places.

#### B.3 Additional file 3

**Filename**: SF3_Zhang_PF_pf_heatshock_PB31_Normal_vs_heatshock

**Format**: CSV

**Description**: Table of the GO terms which are significant before or after correction (exclusively) when comparing PB31 cultured with or without heat shock conditions. Fisher test value before and after correction by BLASE is given.

